# Endogenous but not sensory-driven activity controls migration, morphogenesis and survival of adult-born neurons in the mouse olfactory bulb

**DOI:** 10.1101/2021.04.21.440775

**Authors:** Kaizhen Li, Katherine Figarella, Xin Su, Yury Kovalchuk, Jessika Gorzolka, Jonas J. Neher, Nima Mojtahedi, Nicolas Casadei, Ulrike B. S. Hedrich, Olga Garaschuk

## Abstract

The development and survival of adult-born neurons is believed to be driven by sensory signaling. By genetically manipulating excitability of adult-born cells (via cell-specific overexpression of either Kv1.2 or Kir2.1 K^+^ channels), longitudinal *in vivo* monitoring of their Ca^2+^ signaling and transcriptome analyses, we show that endogenous but not sensory-driven activity governs migration, morphogenesis, survival, and functional integration of adult-born juxtaglomerular neurons in the mouse olfactory bulb. The proper development of these cells required fluctuations of cytosolic Ca^2+^ levels, phosphorylation of CREB, and pCREB-mediated gene expression. Attenuating Ca^2+^ fluctuations via K^+^ channel overexpression strongly downregulated genes involved in neuronal migration, differentiation, and morphogenesis and upregulated apoptosis-related genes, thus locking adult-born cells in the vulnerable and immature state. Together, the data reveal signaling pathways connecting the endogenous intermittent neuronal activity/Ca^2+^ fluctuations as well as proper Kv1.2/Kir2.1 K^+^ channel function to migration, maturation, and survival of adult-born neurons.

## Introduction

The rodent olfactory bulb (OB) is a highly plastic brain region receiving new neurons throughout life. Cumulative evidence points to an important role of these cells for the fine tuning of odor perception/discrimination, facilitation of the task-dependent pattern separation as well as learning and memory (1–6). Generated in the subventricular zone (SVZ) of the lateral ventricle (7, 8), adult-born cells migrate along the rostral migratory stream (RMS) into the OB and differentiate into local GABAergic interneurons: granule cells (GCs) in the granule cell layer and juxtaglomerular cells (JGCs) in the glomerular layer (9) of the bulb. Many molecules including GABA, glutamate, dopamine, serotonin, BDNF, and cAMP response element-binding protein (CREB) influence the adult OB neurogenesis (10–14). Yet, the exact mechanisms underlying the migration, maturation, and incorporation of adult-born neurons into the existent OB circuitry remain unclear. So far the data points to the key role of sensory experience (15–18). Indeed, adult-born cells respond to odorants right after their appearance in the OB (19, 20), develop larger and more complex dendritic trees in the odor-enriched environment (21), and manipulations increasing sensory-driven activity like odor enrichment (22, 23), odor discrimination training (5, 24) or olfactory learning (25) increased the survival and integration of adult-born cells. Conversely, manipulations reducing sensory-driven activity like naris occlusion (16, 26–28), naris cauterization and benzodiazepine treatment (29), knocking-out olfactory receptors in olfactory sensory neurons (30) or their axotomy (31) were reported to decrease survival and integration of adult-born neurons. However, there is also opposing data. Thus, newborn neurons developing in sensory-deprived bulbs had normal dendritic morphology and dynamics (32). Depending on the protocol, olfactory enrichment either did not affect or even decreased survival of adult-born JGCs (33). Besides, the apoptosis of adult-born neurons was induced by the chemogenetic activation of higher order odor-processing brain areas (34), synapsing on these cells. According to the single-cell RNAseq data, odor deprivation increased and olfactory learning decreased numbers of cells in the immature and the majority of more mature adult-born cell clusters (15). On the protein level, the sensory experience-driven changes were even smaller. Still, odor deprivation and not olfactory learning increased the survival of calretinin-positive adult-born GCs (15).

In contrast to adult-born GCs, which migrate straight to their final destinations and integrate therein, adult-born JGCs enter the 3-4 weeks long pre-integration phase (35). During this phase, adult-born JGCs undergo a millimeter-long lateral migration (35), extensively grow and prune their dendritic trees (19, 36) and exhibit ongoing endogenous activity (37). Therefore, in the current study we have tested the role of this endogenous activity for migration, morphogenesis, and survival of adult-born JGCs. To do so, we genetically suppressed their excitability and monitored their developmental history by means of longitudinal *in vivo* two-photon imaging. To understand the molecular pathways involved, we analyzed the transcriptome of adult-born cells right after their arrival into the OB.

## Results

### Alteration of endogenous but not sensory-driven activity impaired the lateral migration of adult-born cells

To suppress the excitability of adult-born cells, we virally overexpressed either outwardly rectifying voltage-gated Kv1.2 channels or inwardly rectifying Kir2.1 channels (38, 39). Bicistronic lentiviruses enabling simultaneous expression of a genetically-encoded Ca^2+^ indicator Twitch-2B and either Kv1.2 or Kir2.1 (Fig, 1A and SI Appendix, Fig. S1A) were injected into the RMS to transduce adult-born cells migrating towards the OB (35). In the control group, mCherry was expressed together with Twitch-2B. The efficiency of virus transduction was confirmed by the level of Twitch-2B fluorescence, the relative amount of the respective mRNA, and the heightened level of Kv1.2 immunofluorescence (SI Appendix, Fig. S1B-F). Ca^2+^ signaling and the developmental history of adult-born JGCs after their arrival in the glomerular layer of the bulb were monitored longitudinally through a chronic cranial window from 8 days post-injection (dpi) till 45 dpi using *in vivo* two-photon imaging.

To analyze the migration of adult-born JGCs, we monitored the cells’ position every 15 min during a 4-hour-long imaging session and compared migration distances of cells, moving during this time window (Fig. 1B). Compared to Twitch-2B/mCherry-expressing control group, the expression of neither Kv1.2 nor Kir2.1 affected the apparent rate of adult-born cells’ arrival in the glomerular layer of the bulb (SI Appendix, Fig. S2). However, it severely impaired cell migration at 8 dpi (Fig. 1C). The median (per mouse) fraction of migrating cells was 33.3 ± 21.7% in the control group but decreased significantly in the Kv1.2 and Kir2.1 groups (12.1 ± 6.6% and 0 ± 5.9%, respectively). Moreover, the median and maximum speed of migrating cells (Fig. 1D, E) as well as the cumulative translocation in 4 hours (Fig. 1F) decreased significantly in the Kv1.2 and Kir2.1 groups compared to the control group. Since adult-born JGCs have a saltatory migration pattern (35), some of them might pause during the 4-hour-long recording period. Therefore, we also evaluated the overall motility of adult-born JGC populations over a 3-day-long time period (8-11 dpi). The cells, which changed their positions between 8 and 11 dpi, were defined as migrating. In the control group, 82.1 ± 8.8% of cells migrated between 8 and 11 dpi (see also (35)). This fraction was significantly lower for the Kv1.2 and Kir2.1 groups (Fig. 1G; 60 ± 21.8% and 54 ± 17% of cells, respectively).

**Figure 1.**
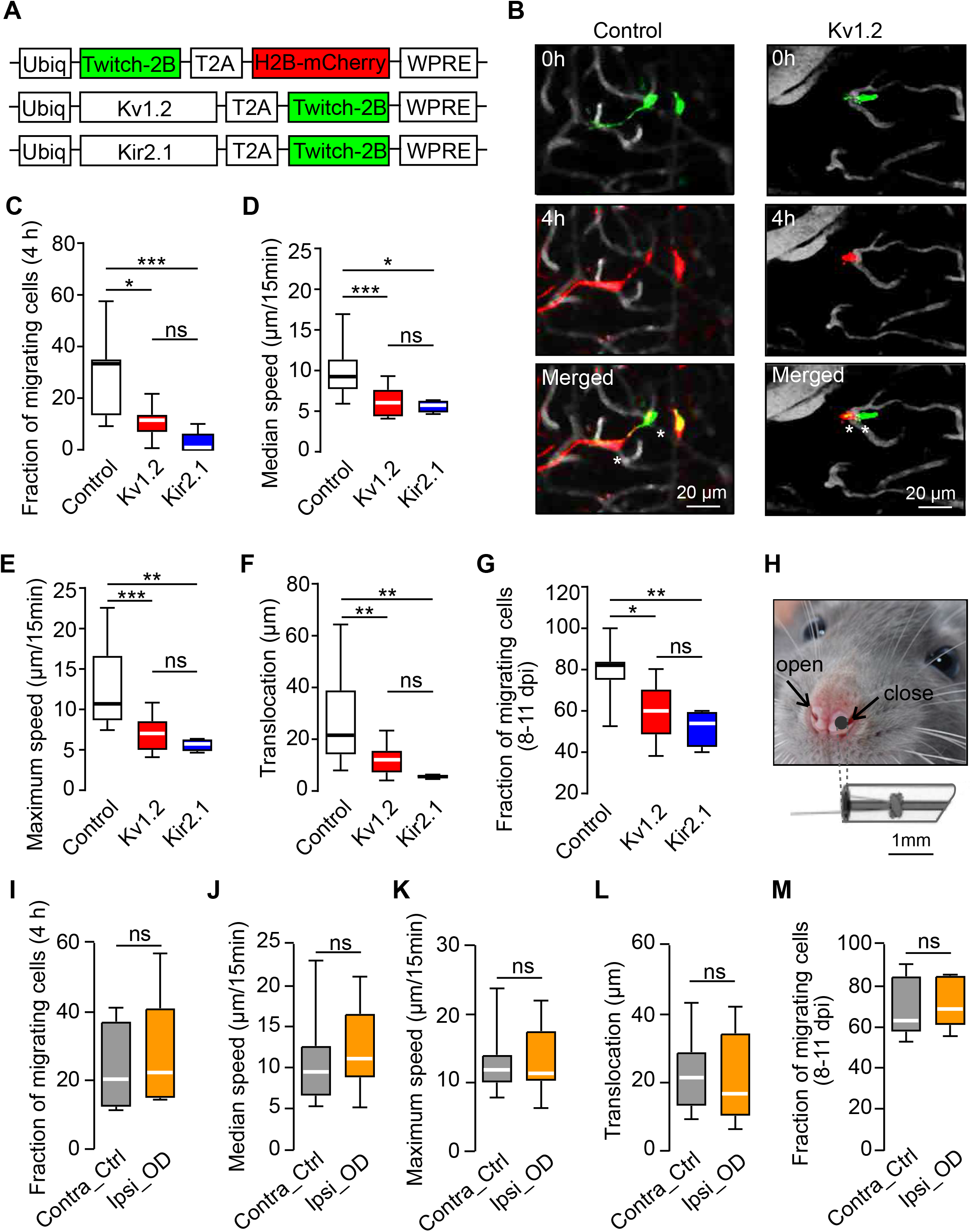
Migration properties of adult-born JGCs from control, Kv1.2, Kir2.1, and odor-deprived groups. (A) Schematics of lentiviral constructs. (B) Pseudocolor images showing migrating control (left) and Kv1.2-expressing (right) adult-born JGCs at different times (timestamps, relative time). Blood vessels are shown in gray, asterisks highlight the positions of migrating cells at 0 and 4h. Data shown in (B-F) and (I-L) are obtained at 8 dpi. (C) Box plot showing the median (per mouse) fractions of migrating JGCs during 4-hour-long recordings in control, Kv1.2, and Kir2.1 groups (n=11, 10 and 6 mice, respectively). (D-F) Box plots showing (per cell) the median (D) or the maximum (E) migration speed and the translocation distance in 4 hours (F); n=29/13, 13/9, 4/6 cells/mice for control, Kv1.2 and Kir2.1 groups, respectively. Three mice had no migrating Kir2.1 cells. (G) Box plot showing the median (per mouse) fractions of JGCs, migrating between 8 and 11 dpi; n=8, 10, 4 mice for control, Kv1.2, and Kir2.1 groups, respectively. (H) Image illustrating the nostril occlusion. (I-M) Box plots showing median (per mouse) fractions of JGCs migrating in 4 hours (I), the median (per cell) (J) and maximum (K) migration speed, the translocation distance in 4 hours (L), and the (per mouse) fraction of cells, migrating between 8 and 11 dpi (M), in contralateral control and ipsilateral odor-deprived hemibulbs (n=16/5 and 19/5 cells/mice for control and odor-deprived groups, respectively). Data are shown as median ± IQR (interquartile range). \**P*<0.05, \*\**P*<0.01, \*\*\**P*<0.001, ns=not significant. Here and below all exact *P* values are listed in SI Appendix, Table S4.

The data suggested that the observed impairment of migration in Kv1.2 and Kir2.1 groups was chronic, as it persisted at a later developmental stage (14 dpi; SI Appendix, Fig. S3). By this time the lateral migration of control adult-born JGCs is known to slow down (35). Likely therefore there was no difference in the fraction of migrating cells as well as the median and maximum migrating speed between control and Kv1.2 groups (SI Appendix, Fig. S3A-C). Yet, the significantly shorter net translocation distance in the Kv1.2 group (SI Appendix, Fig. S3D) suggested that cell motility was impaired. At 14 dpi Kir2.1 expressing cells were not migrating at all (SI Appendix, Fig. S3A). This finding is consistent with the somewhat stronger impact of Kir2.1 on cell migration compared to Kv1.2, visible throughout the data (Fig. 1C-G).

Next, we explored whether modulation of sensory input impacts the migration of adult-born JGCs. Because odors are inhaled through the nostrils into two segregated nasal passages (40), we occluded one nostril and thus also the ipsilateral hemibulb, leaving the other nostril open. Nostril occlusion (Fig. 1H), which started at 5 dpi, completely blocked odor-evoked Ca^2+^ transients in adult-born JGCs residing in the ipsilateral hemibulb (measured at 20 dpi), while leaving the adult-born JGCs in the contralateral control hemibulb unaffected (compare SI Appendix, Fig. S4 to Fig. 4). Surprisingly, odor deprivation was unable to affect any of the migration parameters mentioned above (Fig. 1I-M). All migration parameters were similar between cells residing in the contralateral control and the ipsilateral odor-deprived hemibulbs. Moreover, odor deprivation did not affect the translocation distance in 12 hours at 6.5-14.5 dpi, repeatedly measured by us previously (see Fig. S4 in ref. (35)). Taken together, these data suggest that endogenous but not sensory-driven activity is critical for the lateral migration of adult-born JGCs during the pre-integration phase.

### Endogenous but not sensory-driven activity controls morphogenesis of adult-born JGCs

Immature adult-born cells in the RMS maintain a spindle shape morphology to enable fast migration but adopt a rather complex dendritic structure shortly after reaching the glomerular layer (19, 30, 41). Surprisingly, even at 20 dpi adult-born JGCs in Kv1.2 and Kir2.1 groups displayed remarkably retarded morphology with significantly shorter total dendritic branch length (TDBL) as well as fewer dendritic branches and intersections (Fig. 2A-D). Control adult-born JGCs had a median (per cell) TDBL of 553 ± 630 µm and branch number of 38 ± 50. These values, however, dropped significantly to 75 ± 197 µm / 122 ± 187 µm (TDBL) and 3 ± 6 / 3 ± 16 (branch number) in Kv1.2 and Kir2.1 groups, respectively (Fig. 2B, C). This finding was not an artifact caused by an inferior resolution of thin dendritic processes in *in vivo* measurements, as similar differences were observed in the fixed tissue by means of immunocytochemistry (SI Appendix, Fig. S5). In 50-µm-thick slices, however, the overall length and complexity of the cells were somewhat reduced due to the slicing procedure. The observed impairment of morphogenesis was likely caused by Kv1.2/Kir2.1-mediated reduction of neuronal activity because expressing the non-conducting dominant-negative mutant of the Kir2.1 channel (Kir2.1mut) (42) within the adult-born cells did not inhibit the morphogenesis of cells under study (SI Appendix, Fig. S6). Moreover, we did not observe any changes in the morphology of adult-born JGCs, residing in the control and odor-deprived hemibulbs (Fig. 2E-H).

**Figure 2.**
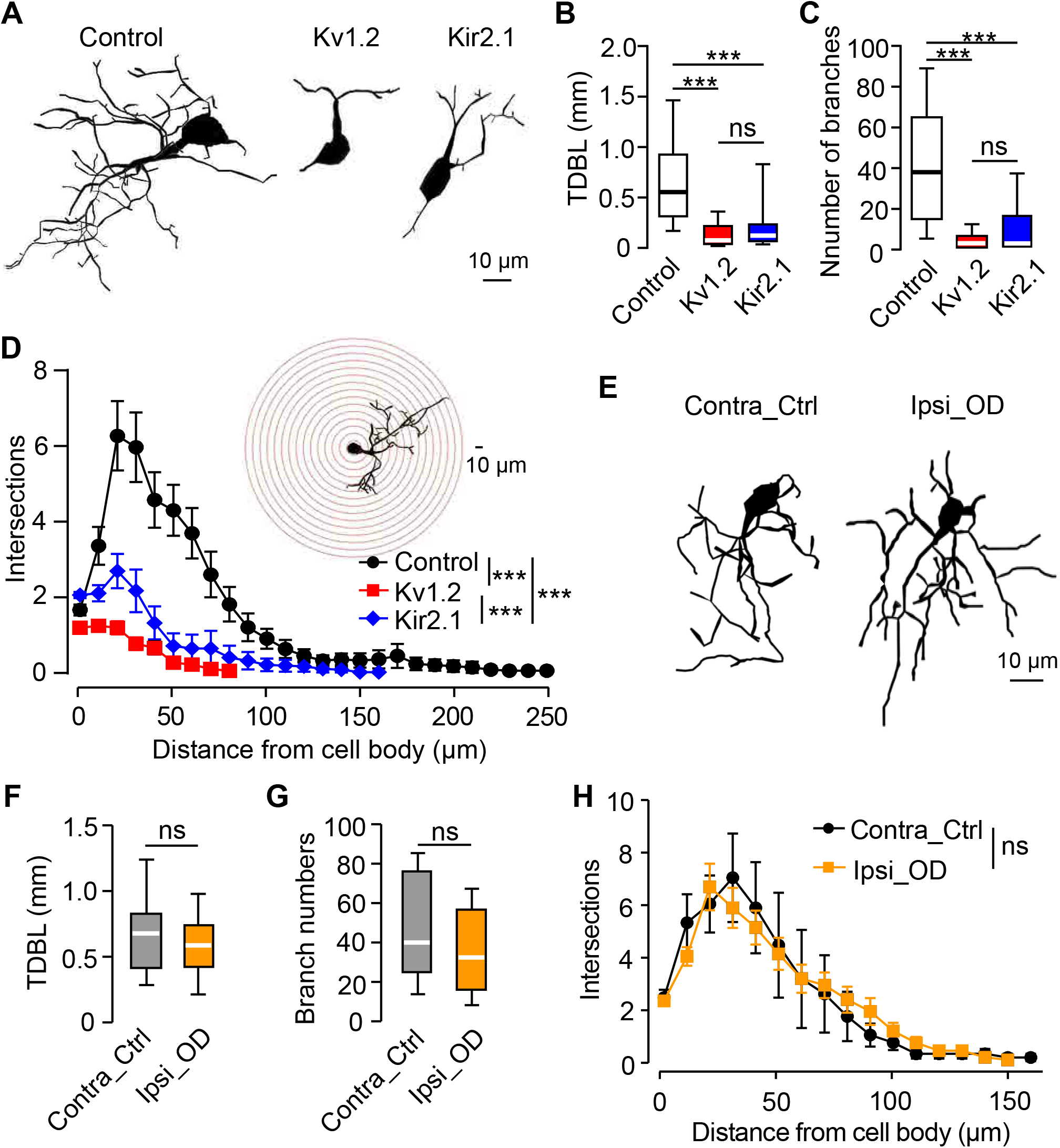
Morphology of adult-born JGCs from control, Kv1.2, Kir2.1, and odor-deprived groups. (A) Sample reconstructions of *in vivo* adult-born JGCs (20 dpi) belonging to control, Kv1.2, and Kir2.1 groups. (B, C) Box plots showing (per cell) the total dendritic branch length (TDBL, (B)) and the number of dendritic branches (C) of adult-born JGCs, imaged *in vivo* at 20 dpi; n=33/7, 36/6, and 46/6 cells/mice for control, Kv1.2, and Kir2.1 groups, respectively. (D) Sholl analysis showing the number of intersections of centered Sholl spheres (10 µm step size, illustrated in the inset) with the dendritic trees of JGCs, belonging to control, Kv1.2 and Kir2.1 groups. n=33/7, 36/6, 46/6 cells/mice for control, Kv1.2 and Kir2.1 groups, respectively. (E) Sample reconstructions of *in vivo* adult-born JGCs (20 dpi) from the contralateral control and ipsilateral odor-deprived hemibulbs. (F, G) Box plots illustrating (per cell) the TDBL (F) and the number of dendritic branches (G) of control and odor-deprived adult-born JGCs. (H) Sholl analysis of the dendritic trees of JGCs from control and odor-deprived hemibulbs; n=7/4 and 20/4 cells/mice for control and odor-deprived groups, respectively. (B) (C) (F) and (G): Data are shown as median ± IQR. (D) and (H): Data are shown as mean ± SEM. \*\*\**P*<0.001, ns=not significant.

Together, these results show that endogenous but not sensory-driven activity is essential for the proper morphological development of adult-born JGCs.

### Alteration of the endogenous activity reduced the survival rate of adult-born JGCs

Next, we analyzed the effect of altered endogenous activity on the survival of adult-born JGCs. Because the rapid migration of adult-born JGCs at the beginning of the pre-integration phase (35) makes the identification of the individual cells difficult, we started to track cells at 14 dpi. We utilized a priori knowledge about the average speed of cell migration (35) to set a safety margin between the analyzed cells and the edge of the field of view (FOV), ensuring that cells, which disappeared from the FOV, did not simply move out. Cells, which stayed at the same position during the recording period, were considered stable surviving cells (Fig. 3A, C). We found that the expression of either Kv1.2 or Kir2.1 significantly reduced the survival rate of adult-born JGCs between 14 and 25 dpi (Fig. 3B). Under these conditions, the mean (per mouse) survival rate was 66 ± 5.6% in control, 40 ± 6.4% in Kv1.2, and 32 ± 5.8% in the Kir2.1 group. Between 25 and 45 dpi, the survival was still inhibited significantly in the Kir2.1 group (35.6 ± 8.9% vs. 82 ± 5.6% in control) but not in the Kv1.2 group (76.3 ± 5.6%) (Fig. 3D), likely because the activity of cells was increasing in the course of development, enabling them to overcome the inhibitory effect of surplus Kv1.2 channels.

**Figure 3.**
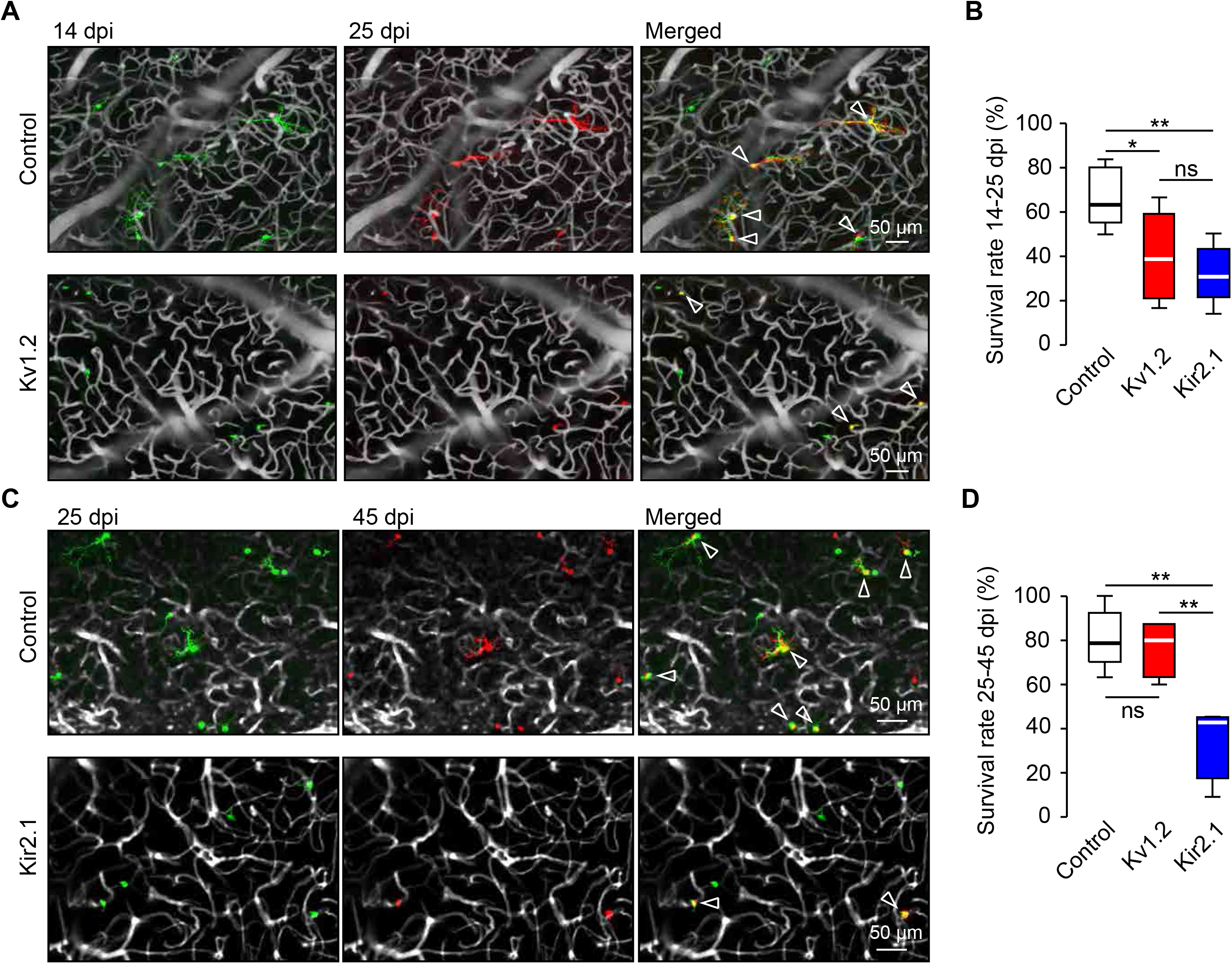
Kv1.2 or Kir2.1 overexpression reduced the survival of adult-born JGCs. (A) Sample maximum intensity projection (MIP) images showing adult-born JGCs at 14 and 25 dpi in control (40-80 µm below the dura) and Kv1.2 (20-50 µm below the dura) groups. Note that in all images the safety margin is removed by cropping, showing just the central part of the image. Arrowheads point to cells, surviving till 25 dpi. (B) Box plot showing the survival rates of adult-born JGCs between 14-25 dpi in control, Kv1.2 and Kir2.1 groups; n=40/5, 81/8 and 52/5 cells/mice for control, Kv1.2 and Kir2.1 groups, respectively. (C) Sample MIP images showing adult-born JGCs at 25 and 45 dpi in control (13-80 µm below the dura) and Kir2.1 (33-65 µm below the dura) groups. Arrowheads point to cells, surviving till 45 dpi. (D) Box plot showing the survival rates of adult-born JGCs between 25-45 dpi in control, Kv1.2, and Kir2.1 groups; n=82/5, 23/5, and 49/4 cells/mice for control, Kv1.2 and Kir2.1 groups, respectively. (B) and (D): Data are shown as median ± IQR. \**P*<0.05, \*\**P*<0.01, ns=not significant.

Taken together, these data suggest that reduced endogenous activity inhibits the survival of adult-born JGCs during the pre-integration phase, with somewhat stronger effects observed in the Kir2.1 compared to the Kv1.2 group.

### Weaker integration of adult-born JGCs with altered endogenous activity into the existent neural circuitry

Next, we explored whether adult-born JGCs belonging to Kv1.2 and Kir2.1 groups were able to integrate into the existent neural network. We used the fraction of odor-responsive adult-born JGCs as well as the amplitude and the area under the curve (AUC) of the odor-evoked Ca^2+^ signals, measured at dpi 20, as a functional readout of the strength of their network integration. We found that the overexpression of either Kv1.2 or Kir2.1 diminished the odor responsiveness of adult-born JGCs (Fig. 4A). The mean (per mouse) fraction of odor responsive cells was 61.5 ± 8.6% in the control group, decreasing slightly (to 51 ± 6.1%) in the Kv1.2 and significantly (to 24.8 ± 6%) in the Kir2.1 groups. The amplitudes and AUCs of odor-evoked Ca^2+^ transients in the Kv1.2 and Kir2.1 groups were significantly smaller than in the control group (Fig. 4B-D). The median (per cell) amplitude of odor-evoked Ca^2+^ transients reached 1.36 ± 1.19 ΔR/R in the control group, 0.83 ± 0.69 ΔR/R in the Kv1.2 group, and 0.85 ± 0.46 ΔR/R in the Kir2.1 group (Fig. 4C). The median (per cell) AUC of odor-evoked Ca^2+^ transients was 5.7 ± 6.3 ΔR/R*s in the control, 3.1 ± 3.2 ΔR/R*s in the Kv1.2 and 2.0 ± 1.7 ΔR/R*s in the Kir2.1 group (Fig. 4D).

**Figure 4.**
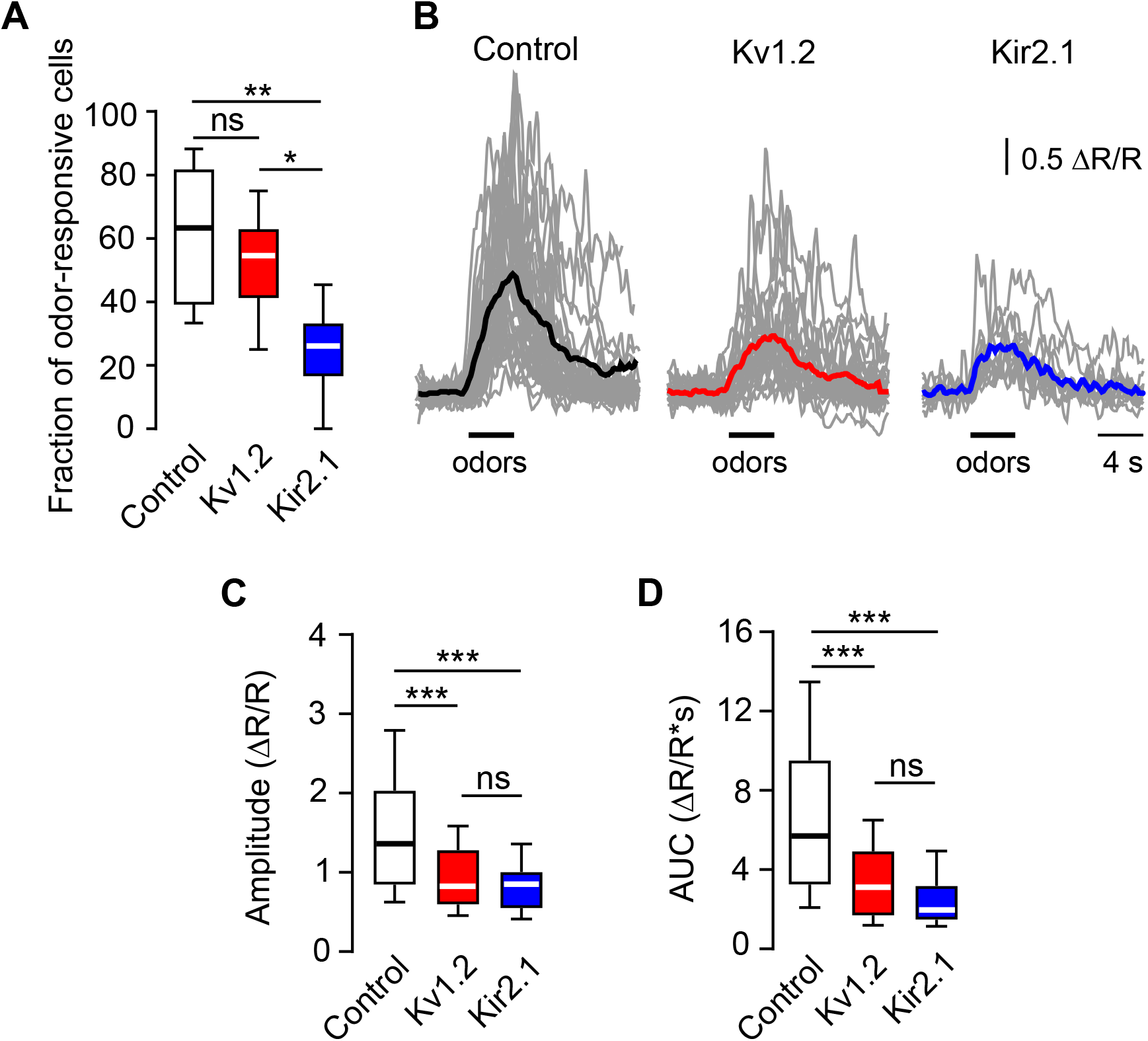
Kv1.2 or Kir2.1 overexpression inhibited odor-evoked Ca^2+^ transients in adult-born JGCs. (A) Box plot showing the fraction of odor-responsive adult-born JGCs in control, Kv1.2 and Kir2.1 groups (20 dpi); n=78/6, 59/7 and 63/6 cells/mice for control, Kv1.2 and Kir2.1 groups, respectively. (B) Sample traces showing the odor-evoked Ca^2+^ transients of adult-born JGCs at 20 dpi. Colored bold traces are the means of all recordings (shown in gray). (C) and (D) Box plots summarizing the median amplitude (C) and area under the curve (AUC, (D)) per cell of odor-evoked Ca^2+^ transients. (B-D) n=82/7, 53/7, and 17/5 cells/mice for control, Kv1.2, and Kir2.1 groups, respectively. Data are shown as median ± IQR (interquartile range). \**P*<0.05, \*\**P*<0.01, \*\*\**P*<0.001, ns=not significant.

Taken together, these data suggest that adult-born JGCs with altered endogenous activity have difficulties to integrate into the existent OB circuitry.

### Spontaneous Ca^2+^ transients in adult-born JGCs were dampened by overexpression of Kv1.2 or Kir2.1 but maintained during odor deprivation

To extract features of the endogenous activity crucial for migration, morphogenesis, survival, and integration of adult-born JGCs, we conducted (at 12 dpi) comparative analyses of the pattern of endogenous activity recorded in control, Kv1.2, Kir2.1 groups as well as in two groups from odor-deprived mice (cells residing in odor-deprived ipsilateral and control contralateral hemibulbs). In adult-born JGCs, the overexpression of Kv1.2 or Kir2.1 did not affect the basal Twitch-2B ratios (reflecting basal levels of the intracellular free Ca^2+^ concentration ([Ca^2+^]_i_)), maximum Twitch-2B ratios, and normalized AUCs (SI Appendix, Fig. S7A-C). However, it profoundly reduced the fraction of cells with spontaneous fluctuations in [Ca^2+^]_i_ (Fig. 5A). Using the Gaussian mixture model (43), 36 out of 60 cells with fluctuations in [Ca^2+^]_i_ (i.e., active cells) were identified in the control group, whereas in the Kv1.2 group only 23 out of 68 cells and in the Kir2.1 group only 18 out of 66 cells were active (Fig. 5B). We also calculated the mean (per mouse) fraction of active cells, amounting to 57.5 ± 7.8% in the control, 33.5 ± 3.1% in the Kv1.2, and 23.5 ± 6.7% in the Kir2.1 group (Fig. 5C). These results demonstrate that the overexpression of Kv1.2 or Kir2.1 significantly dampens spontaneous fluctuations in [Ca^2+^]_i_ in adult-born JGCs.

**Figure 5.**
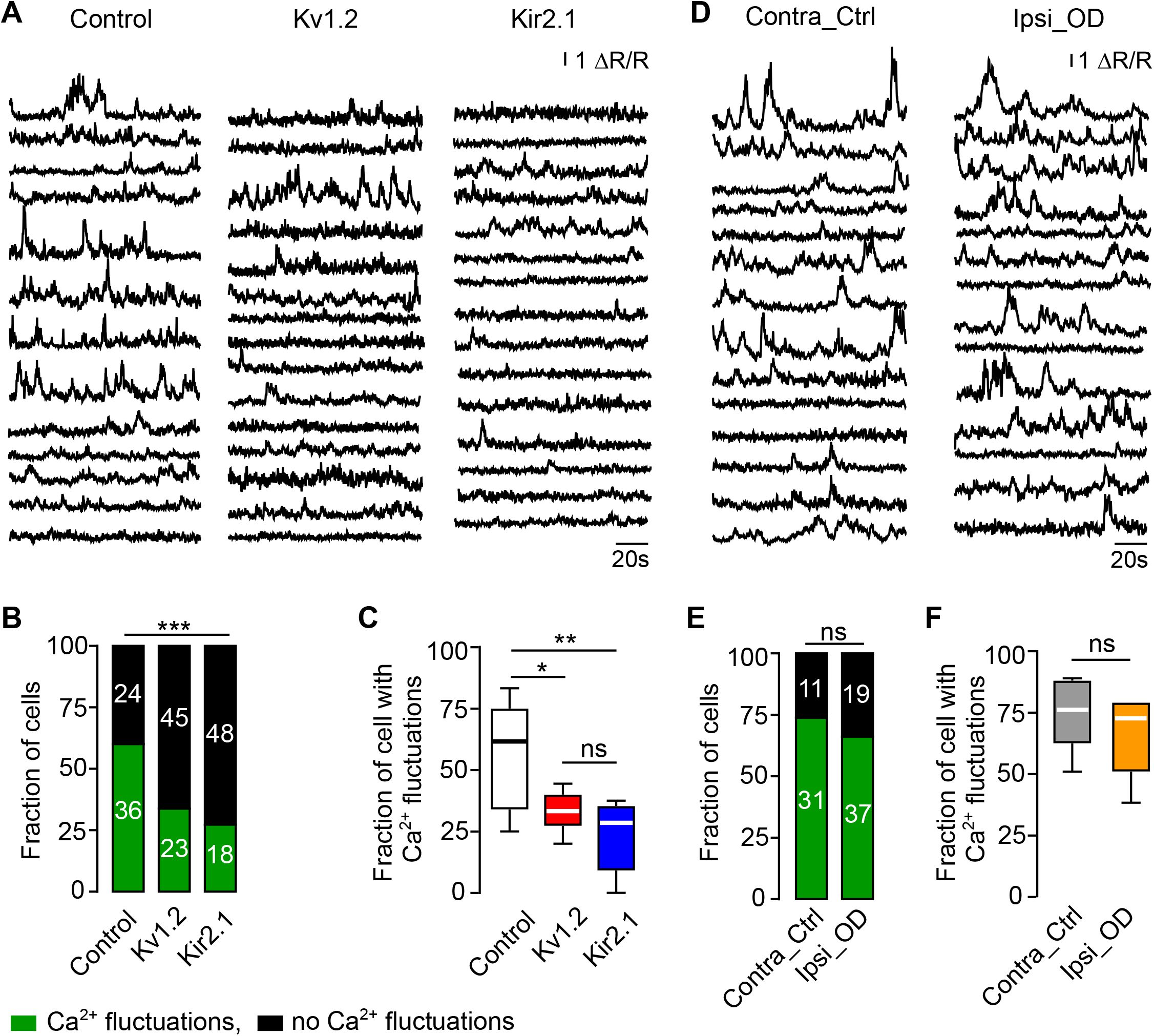
Kv1.2 or Kir2.1 overexpression perturbed the pattern of spontaneous Ca^2+^ transients in adult-born JGCs while odor deprivation did not. (A) Sample traces illustrating spontaneous Ca^2+^ transients obtained at 12 dpi from adult-born JGCs in control, Kv1.2, and Kir2.1 groups in awake mice. (B) Stacked bars showing the fractions of cells with and without fluctuations in [Ca^2+^]_i_. Here and in (E) the numbers of cells in each category are given within the bars. (C) Box plot illustrating the median (per mouse) fractions of adult-born JGCs with fluctuations in [Ca^2+^]_i_. (B, C) n=60/8, 68/7, and 66/5 cells/mice for control, Kv1.2, and Kir2.1 groups, respectively. (D) Sample traces illustrating spontaneous Ca^2+^ transients of adult-born JGCs from contralateral control and ipsilateral odor-deprived hemibulbs. (E) Stacked bars showing the fractions of cells with and without fluctuations in [Ca^2+^]_i_ in control and odor-deprived groups. (F) Box plot illustrating the median (per mouse) fractions of adult-born JGCs with fluctuations in [Ca^2+^]_i_ in control and odor-deprived groups. (E, F) n=34/4 and 47/4 cells/mice for contralateral control and ipsilateral odor-deprived groups, respectively. (C) and (F): Data are shown as median ± IQR (interquartile range). \**P*<0.05, \*\**P*<0.01, ns=not significant.

In contrast, no significant changes in the pattern of endogenous activity were found in odor-deprived cells (Fig. 5D, E, F). The mean fraction of active cells per mouse was 74.2 ± 6.7% in the contralateral control and 65.4 ± 7.5% in the ipsilateral odor-deprived group (Fig. 5F). Both values were similar to those measured in the control group. Besides, there was no significant difference in the basal and maximum Twitch-2B ratios as well as AUCs between the contralateral control and the ipsilateral odor-deprived groups (SI Appendix, Fig. S7D-F). However, we noticed that in both the contralateral control and the ipsilateral odor-deprived groups the basal Twitch-2B ratios and AUCs were slightly but significantly smaller than the respective values measured in mice without nostril occlusion (i.e., control, Kv1.2, and Kir2.1 groups; Kruskal-Wallis test: for basal Twitch-2B ratios, *P* = 5.1×10^-5^; for AUCs, *P* = 5.2×10^-5^). As migration and morphology were not affected in odor-deprived mice, these data suggest that these parameters were not important for the proper development of adult-born JGCs.

Together, these data identify the patterned endogenous activity consisting of fluctuations in [Ca^2+^]_i_ as a key parameter governing the migration, morphogenesis and survival of adult-born JGCs.

### The role of pCREB for regulation of early neuronal development

Phosphorylation of the cAMP response element binding protein (CREB) at the Ser133 residue is known to play an important role in the differentiation and survival of adult-born cells in the OB (11, 44). Typically, pCREB is abundant in developing and low in mature neurons (45). We, therefore, measured the levels of pCREB in our 5 experimental groups (control, Kv1.2, Kir2.1, contralateral control, ipsilateral odor-deprived). Adult-born JGCs in the Kv1.2 and Kir2.1 groups showed a significant decrease in the level of pCREB both during the early (10 dpi, Fig. 6A, B) and the late (28 dpi, Fig. 6C) pre-integration phase. At 10 dpi, the mean (per mouse) relative pCREB level (i.e., normalized to the mean fluorescence intensity of pCREB of all surrounding mature NeuN-positive cells) of adult-born JGCs was 19.7 ± 1.7 in control but decreased sharply to 5.6 ± 0.7 in the Kv1.2 and 5.8 ± 0.8 in the Kir2.1 group. The pCREB levels in the odor-deprived mice were similar to that found in the control group, amounting to 16.8 ± 2.8 in the contralateral control and 17.9 ± 2.5 in the ipsilateral odor-deprived group and significantly different from that found in the Kv1.2/Kir2.1 groups (Fig. 6B). As the pCREB level in adult-born cells gradually decreases during their maturation (45), at 28 dpi the respective values were 4.6 ± 0.5 for the control, 1.5 ± 0.2 for the Kv1.2, and 1.4 ± 0.2 for the Kir2.1 group (Fig. 6C).

**Figure 6.**
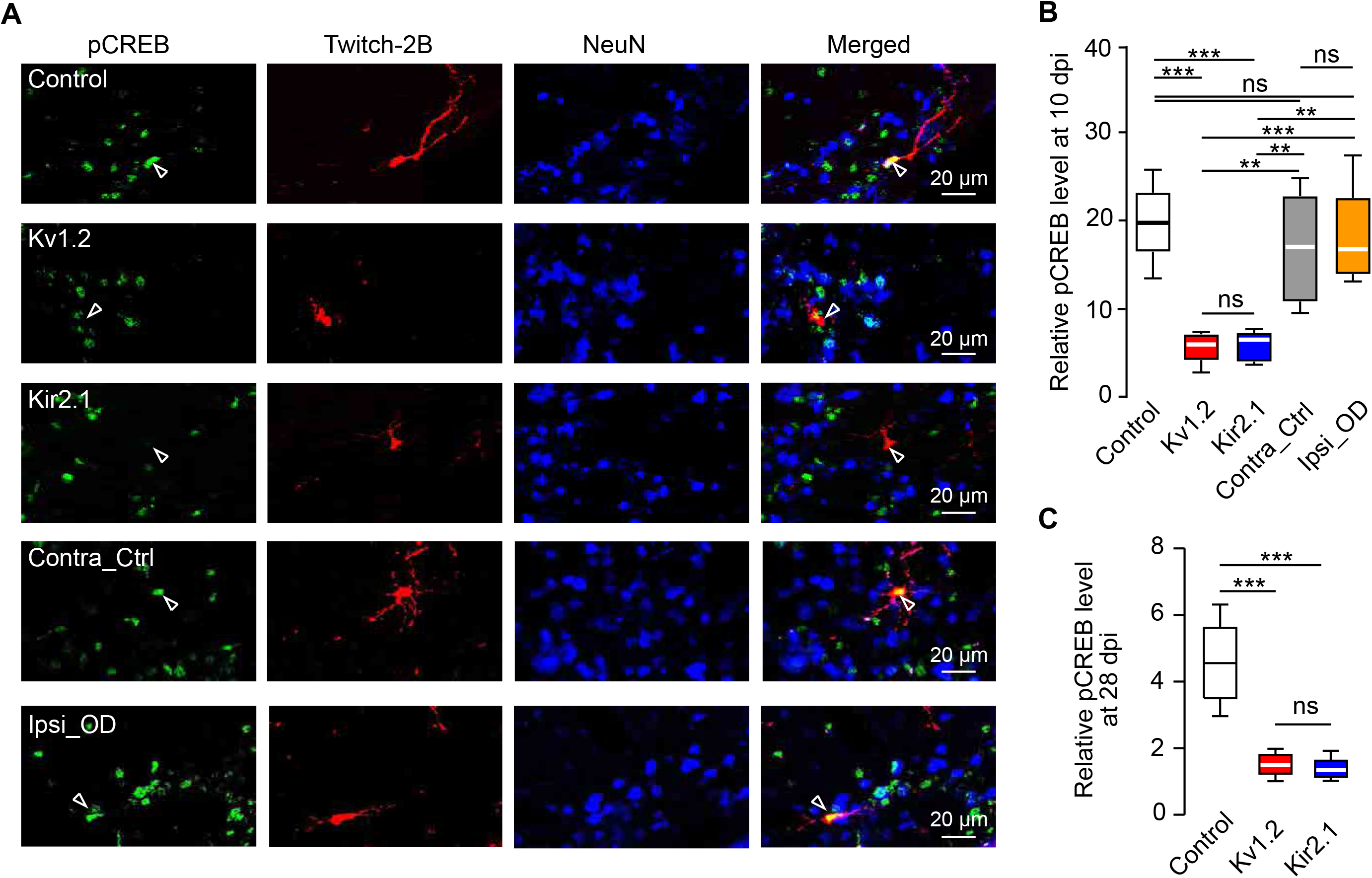
Kv1.2 or Kir2.1 overexpression downregulated pCREB signaling pathway. (A) Sample MIP images (20 µm depth) showing pCREB-, Twitch-2B- and NeuN-positive cells in the glomerular layer of the OB slices from control, Kv1.2, Kir2.1, Contra_Ctrl, and Ipsi_OD groups at 10 dpi. Twitch-2B labels the adult-born JGCs whereas NeuN labels the mature neurons. Arrowheads highlight the location of adult-born JGCs. (B, C) Box plots illustrating the median (per mouse) relative pCREB levels of adult-born JGCs in different experimental groups, as indicated, at 10 dpi (B) and 28 dpi (C). (B) n=152/6, 230/6, 177/ 5, 129/5, and 94/5 cells/mice for control, Kv1.2, Kir2.1, Contra_Ctrl, and Ipsi_OD groups, respectively. (C) n=110/5, 143/5, and 161/5 cells/mice for control, Kv1.2, and Kir2.1 groups, respectively. Data are shown as median ± IQR (interquartile range). \*\**P*<0.01, \*\*\**P*<0.001, ns=not significant.

These data show that in adult-born JGCs with dampened endogenous activity, the pCREB signaling pathway is strongly inhibited throughout the pre-integration phase.

### Transcriptomic analyses of pathways linked to pCREB signaling

To investigate the molecular pathways that might link the dysregulation of JGCs’ Ca^2+^ signaling to the reduction in pCREB, we isolated adult-born cells belonging to the control, Kv1.2, and Kir2.1 groups using fluorescence-activated cell sorting (FACS) and determined their transcriptomic profiles using next-generation sequencing (Fig. 7). For a subset of genes (i.e., one representative member of pathways analyzed in Fig. 7), the data obtained were further validated by quantitative real-time PCR (SI Appendix, Fig. S8). Because the FACS-sorted cells, isolated from the OB, also contained adult-born GCs, we ensured that adult-born GCs recapitulated the impairment in morphology and CREB phosphorylation, described above for adult-born JGCs (SI Appendix, Fig. S9).

**Figure 7.**
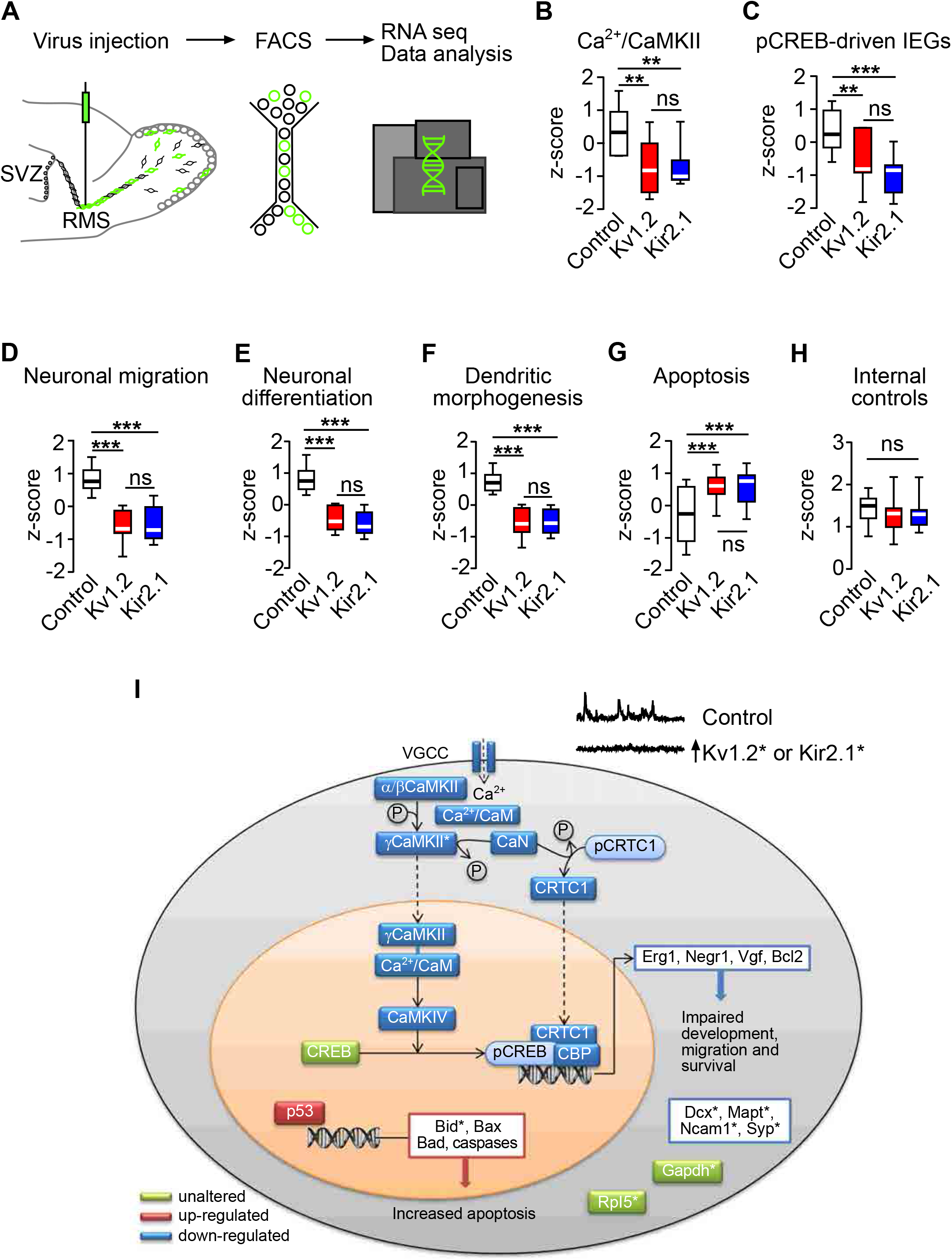
Transcriptome analyses of adult-born cells in control, Kv1.2, and Kir2.1 groups. (A) Schematic illustration of the workflow used for RNAseq. (B-H) Box plots showing the expression levels of genes in the modules of Ca^2+^/CaMKII (B), pCREB-driven IEGs (C), neuronal migration GO:0001764 (D), neuronal differentiation GO:0045664 (E), dendritic morphogenesis GO:0048813 (F), apoptotic process GO:0006915 (G), and 9 housekeeping genes serving as internal controls (H). All transcripts included in the box plots are listed in SI Appendix, Table S1. n = 2 biological replicates (12 mice in total, 2 mice per replicate per group). (I) The schematic summary of the genes/pathways involved. See the discussion for further details. Asterisks indicate genes, expression of which was verified by qPCR. Data are shown as median ± IQR (interquartile range). \*\**P*<0.01, \*\*\**P*<0.001, ns=not significant.

As expected, the data revealed a significant upregulation of the *Kcna2* and *Kcnj2* mRNAs in the Kv1.2 and Kir2.1 groups, respectively (SI Appendix, Fig. S1E, F), as well as a stable expression pattern of housekeeping genes (Fig. 7H, SI Appendix, Fig. S8G). Interestingly, the cells in the Kv1.2 and Kir2.1 groups showed a significantly lower expression of genes participating in the transport of the Ca^2+^/Calmodulin complex (Ca^2+^/CaM) into the nucleus, such as several isoforms of the Ca^2+^/CaM-dependent protein kinase II (CaMKII) (Fig. 7B; for details, see SI Appendix, Table S1). Activation of this shuttle pathway is mediated by Ca^2+^ influx through either voltage-gated L-type Ca^2+^ channels (VGCC; Fig. 7I) or NMDA receptors (Fig. S10A). The final step in the cascade involves activation of the CaMKIV, which is responsible for pCREB phosphorylation. Transcripts, encoding the L-type Ca^2+^ channels, NMDA receptors, and CaMKIV, were also downregulated in the Kv1.2 and Kir2.1 groups (Fig, 7B, I and SI Appendix, Fig. S10A). Moreover, additional molecules, required to fully activate the transcription complex of pCREB, like the CREB-regulated transcription coactivator 1 (CRTC1) and calcineurin (CaN), which dephosphorylates CRTC1 to allow its translocation into the nucleus and binding to the transcription unit, were also downregulated (Fig. 7I). Next, we investigated further molecules, which act as coactivators or repressors within the transcription unit of pCREB, e.g., LIM domain only 4 (LMO4), DRE-Antagonist Modulator (DREAM), Calcium-Responsive Transactivator (CREST)(46, 47). The expression profile of these molecules was consistent with the decreased transcriptional activity of pCREB (SI Appendix, Table S2). Of note, the other major pathway leading to CREB phosphorylation such as the MAPK/ERK pathway remained unchanged (SI Appendix, Fig. S10B).

In accordance with the above findings, we observed a decreased expression of pCREB-driven immediate early genes (e.g., early growth response 1 (*Egr1*), neuronal growth regulator 1 (*Negr1*), etc.; Fig. 7C, I) and downregulation of pathways responsible for neuronal migration (GO:0001764), differentiation (GO:0045664) as well as dendritic morphogenesis (GO:0048813) in the Kv1.2 and Kir2.1 groups (Fig. 7D-F, I; SI Appendix, Fig. S8, Table S1). These results are consistent with the *in vivo* functional properties of adult-born JGCs overexpressing Kv1.2 or Kir2.1 channels. In line with the reduced survival of adult-born JGCs in the Kv1.2 and Kir2.1 groups (Fig. 3), the transcripts involved in the apoptotic process (GO:0006915) were upregulated, while the anti-apoptotic genes, like the apoptosis regulator Bcl2, were downregulated (Fig. 7G, I; SI Appendix, Fig. S8, Table S1).

Taken together, these data identify a defined signaling pathway, connecting the specific pattern of endogenous neuronal activity to downstream signaling cascades regulating migration, differentiation, morphogenesis, and survival of adult-born neurons.

## Discussion

The current study examined the role of endogenous as well as sensory-driven activity for the key aspects of adult-born JGCs’ development, including migration, differentiation, morphogenesis, and survival, and identified the possible molecular mechanisms involved. Surprisingly, and against the widespread belief in the field (see Introduction), sensory-driven activity had little impact on the migration and morphogenesis of adult-born JGCs. Instead, the proper development of adult-born JGCs critically depended on the endogenous fluctuations in [Ca^2+^]_i_, phosphorylation of CREB, and likely also the CRTC1 translocation to the nucleus (Fig. 7I). Interestingly, the differences in the steady-state levels of [Ca^2+^]_i_, reflected in the basal Twitch-2B ratios, turned out to be much less important. Moreover, our transcriptomic data identified the candidate genes, which relay this Ca^2+^/pCREB/CRTC1 signal further to govern the integration of these newly-generated cells into existing neural circuitry.

According to our current as well as previous (35) data, sensory-driven activity had surprisingly little impact on migration, differentiation, and morphogenesis of adult-born JGCs, despite their vivid odor responsiveness already at dpi 9 (19). Consistently, we did not observe any decrease in CREB phosphorylation in odor-deprived hemibulbs. Regarding morphogenesis, our data are consistent with that of A. Mizrahi (32) but fits less well to his later data showing almost doubling of the TDBL in cells, exposed to an odor-enriched environment (21). The data on OB GCs provides a similarly inconsistent picture. On the one hand, the migration and morphology of adult-born GCs were normal in anosmic mice lacking any electrical activity in the olfactory epithelium (30), as was the GCs morphology in young adult mice after 5 weeks of naris occlusion (48). On the other hand, 30-40 days of naris occlusion starting at postnatal days 4-5 caused a significant reduction of TDBL of GCs (49). Moreover, naris occlusion also impeded the migration of neuroblasts along the RMS and the tenascin-R-dependent radial migration of neuroblasts from the RMS to the olfactory bulb (16, 17). Longitudinal monitoring of cells’ basal and manipulation-induced activity (e.g., by means of Ca^2+^ imaging (36)), as well as a clear separation between an early postnatal and adult neurogenesis, might help to solve the above discrepancies in the future. The literature data, however, are consistent regarding the negative impact of naris occlusion on the survival and synapse/spine density of adult-born OB interneurons (27-30, 48, 50). Here, we show that the survival of adult-born JGCs is significantly impaired by blocking endogenous activity (Fig. 3). Whether the sensory deprivation-induced neuronal loss (28) is additive, remains to be explored.

The endogenous activity is present during all stages of the adult OB neurogenesis, i.e., in neural progenitor cells in the SVZ (51), in neuroblasts in the RMS (12, 52) and adult-born cells in the subependymal (53) as well as glomerular (37) layers of the OB. The need for the intermittent activity or ongoing fluctuations in [Ca^2+^]_i_ for the maturation process can be explained by the interplay of the CaMKII- and the CRTC1-dependent pathways. Both pathways are triggered by the activation of L-type voltage-gated Ca^2+^ channels (and NMDA receptors) and require the Ca^2+^/calmodulin (CaM) complex (46, 54, 55). While the CaMKII-dependent pathway leads to the γCaMKII-mediated shuttling of Ca^2+^/CaM to the nucleus and the subsequent phosphorylation of CREB (46, 54, 56), the calcineurin (CaN)-mediated dephosphorylation of CRTC1 (also known as the transducer of regulated CREB 1 (TORC1)) supports its translocation to the nucleus, where it is required for the pCREB-driven gene expression (57, 58). One of such pCREB/CRTC1-dependent target genes is salt-inducible kinase 1 (SIK1), which phosphorylates CRTC1, triggering its export from the nucleus and thus arresting pCREB/CRTC1-mediated transcription (57). Via this mechanism long-lasting steady-state elevation of [Ca^2+^]_i_ weakens rather than potentiates the pCREB-mediated gene expression. Moreover, in the Kv1.2 and Kir2.1 groups analyzed in our study, we also observed the downregulation of transcripts, encoding the proteins involved in these important pathways (Fig. 7), thus further impairing the signal transduction from the cell membrane to the nucleus. Interestingly, the molecular pathways described above are similar to the ones governing the activity-dependent synaptic plasticity as well as memory acquisition and retrieval in the adult brain (46, 59, 60).

To modify the endogenous neuronal activity and accompanying Ca^2+^ signaling, we overexpressed either Kv1.2 or Kir2.1 K^+^ channels, thus mimicking their gain of function. Interestingly, gain-of-function mutations of both Kv1.2 and Kir2.1 channels are indeed found in clinical studies and are often accompanied by either developmental epileptic encephalopathies or somewhat “milder” symptoms like autism, ataxia, brain atrophy, and myoclonic seizures (38, 61–63). Strikingly, the gain-of-function mutations of Kv1.2 channels, which are supposed to promote neuronal repolarization and termination of neuronal firing, caused more severe symptoms in terms of epilepsy, ataxia, and intellectual disability than the loss-of-function mutations, which are supposed to promote neuronal hyperactivity (61, 62). By showing that the “gain-of-function” of either Kv1.2 or Kir2.1 channels impairs migration, differentiation, and morphogenesis of the developing inhibitory neurons, our data provide a plausible explanation for these counterintuitive findings. Moreover, our findings strongly suggest that adult and neonatal neurogenesis share common molecular pathways (39, 57, 64–69). Therefore, we envisage the retarded interneuron development as a key underlying mechanism of the aforementioned developmental pathologies. These pathologies are likely further exacerbated by increased apoptosis of the interneuronal population. According to the literature, apoptosis can be directly triggered by the enhanced transmembrane K^+^ efflux (reviewed in ref. (70)). The decreased expression of anti-apoptotic (*Bcl2*) and the increased expression of pro-apoptotic (*Bid, Bax, Bad, caspases*) genes, revealed by our study, is consistent with our longitudinal imaging data, documenting decreased survival of adult-born cells, and provides a second hypothesis for the gain-of-function-mediated epileptic encephalopathy in clinical settings. Besides, the mice from Kv1.2/Kir2.1 groups are expected to have some deficits in the fine tuning of odor perception/discrimination, facilitation of the task-dependent pattern separation as well as learning and memory, consistent with the aforementioned role of adult-born cells in the OB.

Taken together, our data show that despite developing in the rich sensory environment and being able to respond to sensory stimuli from early on (19, 20, 71), adult-born neurons strongly rely on cell-intrinsic activity for their migration and morphogenesis, being in this respect largely similar to neonatal neurons. By providing several missing links, our data revealed stringent signaling pathways connecting the intermittent neuronal activity/ongoing fluctuations in [Ca^2+^]_i_ on one side and the increased transmembrane K^+^ efflux on the other side to neuronal migration, maturation, and survival.

## Acknowledgments

We thank E. Zirdum, A. Weible and K. Schöntag for technical assistance, C. Lois for providing Mgi-Kir2.1wt and Mgi-Kir2.1mut plasmids, and O. Rieß, H. Lerche, Y. Liang, S. Ciocchi, and M. Sakaguchi for comments on the manuscript. This work was supported by the DFG grants GA 654/14-1 to O.G and GA 654/13-1/HE8155/1-1 (belonging to the Research Unit FOR-2715) to O.G. and U.H.-K. The NCCT is financed by the DFG grant INST 37/1049-1.

## Author Contributions

O.G. conceived the study, K.L., K.F., X.S., Y.K., J.J.N., N.M., N.C., U.H.-K. performed experiments and/or data analyses. K.L., K.F., and O.G. wrote and all authors approved the manuscript.

---

**Figure S1.**
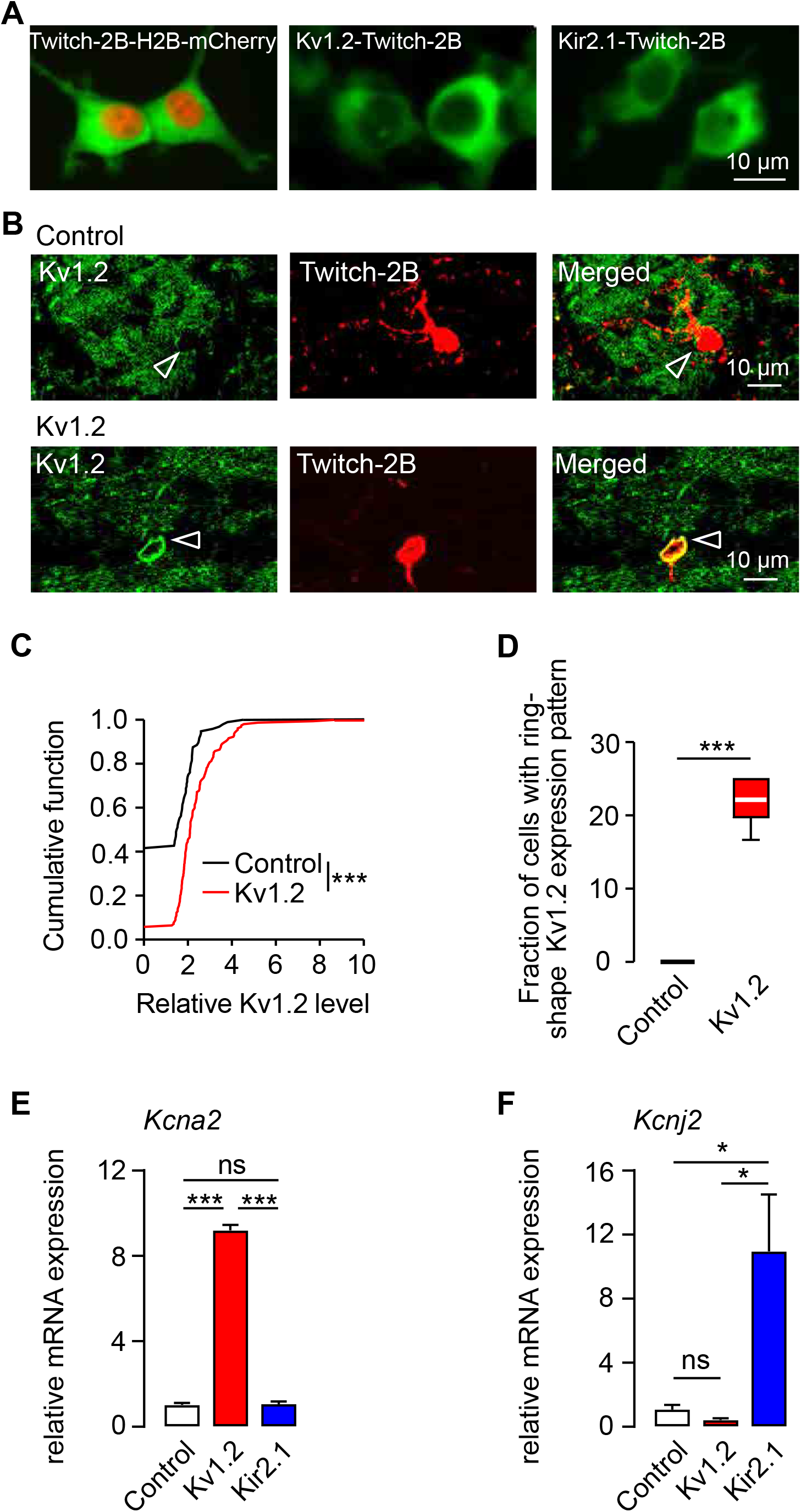
related to Figure 1. Bicistronic lentiviral vectors induced simultaneous expression of Kv1.2/Kir2.1 and Twitch-2B. (A) Sample images showing the transient expression of the lentiviral vectors in HEK-293T cells. (B) Sample MIP immunofluorescence images (20 µm depth) showing the expression of Kv1.2 in adult-born JGCs at 10 dpi in control (upper panel) and Kv1.2 (lower panel) groups. Arrowheads highlight the location of adult-born JGCs. (C) Cumulative distributions of the relative Kv1.2 expression levels in control and Kv1.2-expressing adult-born JGCs. (D) Box plot showing the median fractions (per mouse) of adult-born JGCs with a ring-shape Kv1.2 expression pattern, likely reflecting enhanced somatic targeting of the protein. (C, D) n=96/6 and 155/6 cells/mice for control and Kv1.2 groups, respectively. (E, F) Bar graphs showing the mean relative expression levels of mRNA encoding for the potassium voltage-gated channel subfamily A member 2 (*Kcna2*, encoding Kv1.2. (E)) and the potassium inwardly-rectifying channel subfamily J member 2 (*Kcnj2*, encoding Kir2.1. (F)) as determined by qPCR analyses of FACS-sorted adult-born cells (see Methods). Data are shown as median ± IQR. \**P*<0.05, \*\*\**P*<0.001, ns=not significant.

**Figure S2.**
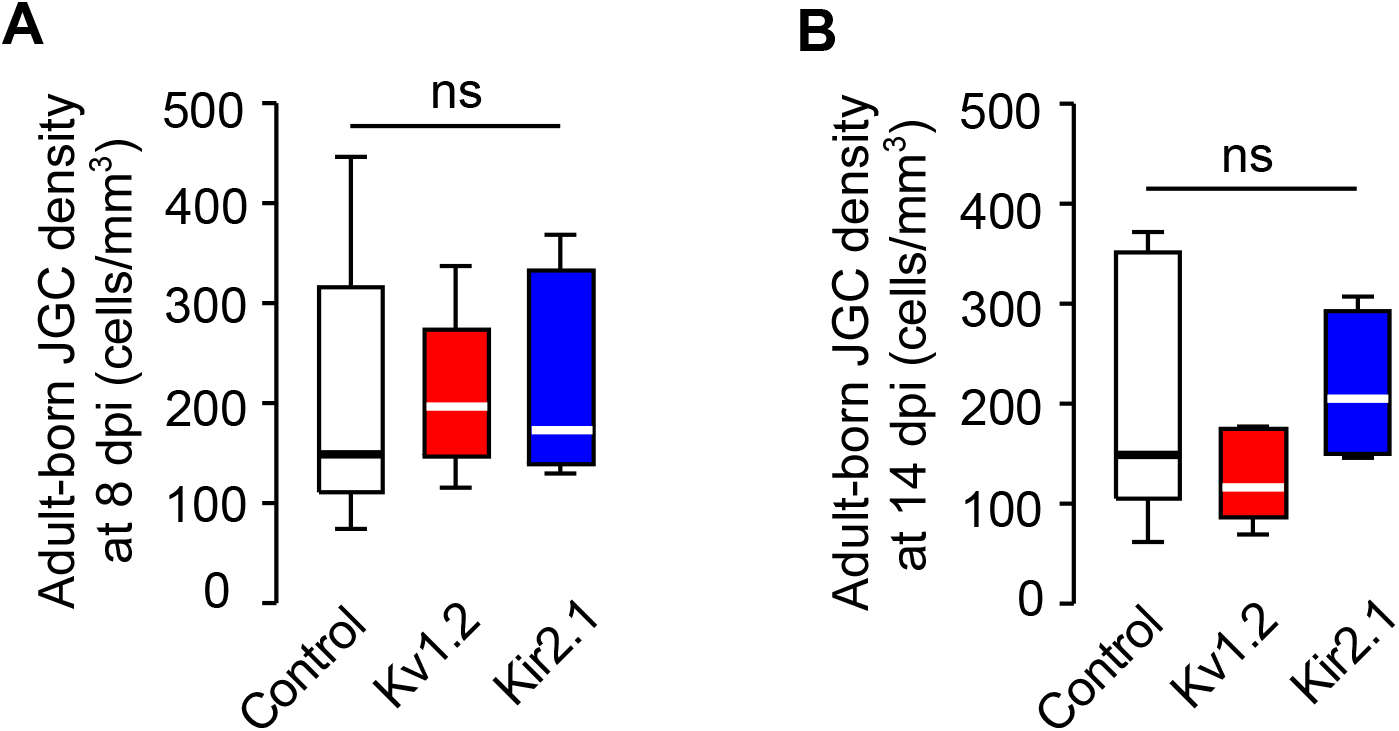
related to Figure1. Kv1.2 or Kir2.1 overexpression did not influence the arrival of adult-born JGCs in the glomerular layer. Box plots showing the median (per 3D stack volume) density of adult-born JGCs in the OB glomerular layer (i.e., number of adult-born JGCs divided by the respective 3D stack volume) at 8 dpi (A) and 14 dpi (B) in control, Kv1.2 and Kir2.1 groups. (A) n = 57/5, 131/9, and 65/6 cells/mice for control, Kv1.2, and Kir2.1 groups, respectively. (B) n = 71/5, 89/7, and 60/4 mice/mice for control, Kv1.2, and Kir2.1 groups, respectively. Data are shown as median ± IQR. ns=not significant.

**Figure S3.**
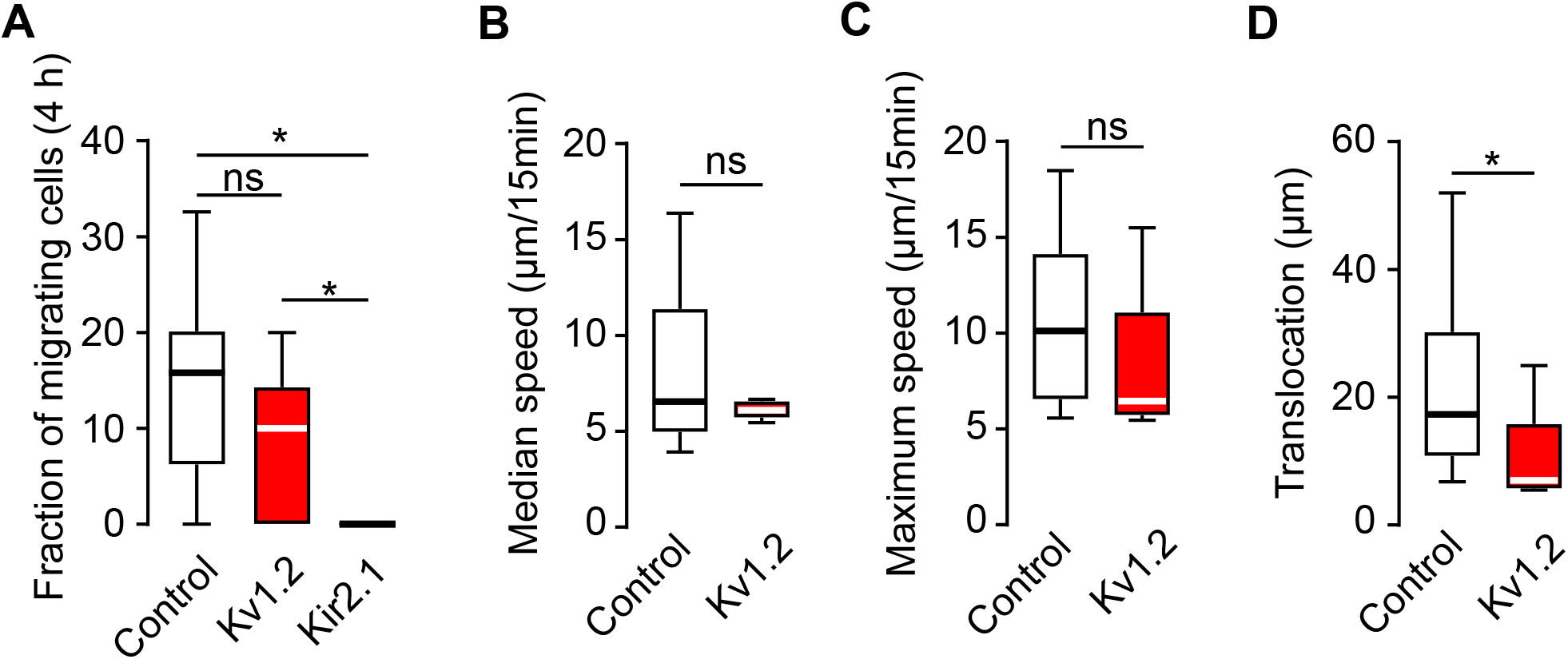
related to Figure1. Kv1.2 or Kir2.1 overexpression impaired the migration of adult-born JGCs at 14 dpi. (A) Box plot showing the median (per mouse) fractions of adult-born JGCs migrating during 4-hour-long recordings in control, Kv1.2, and Kir2.1 groups. Here and below n=30/10, 5/7, and 0/4 migrating cells/mice for control, Kv1.2 and Kir2.1 groups, respectively. In total, we analyzed 216/10, 86/7, and 60/4 cells/mice for control, Kv1.2, and Kir2.1 groups, respectively. (B-D) Box plots showing (per cell) the median migration speed (B), maximum migration speed (C), and the translocation distance in 4 hours (D) of adult-born JGCs in control and Kv1.2 groups. Data are shown as median ± IQR. \**P*<0.05, ns=not significant.

**Figure S4.**
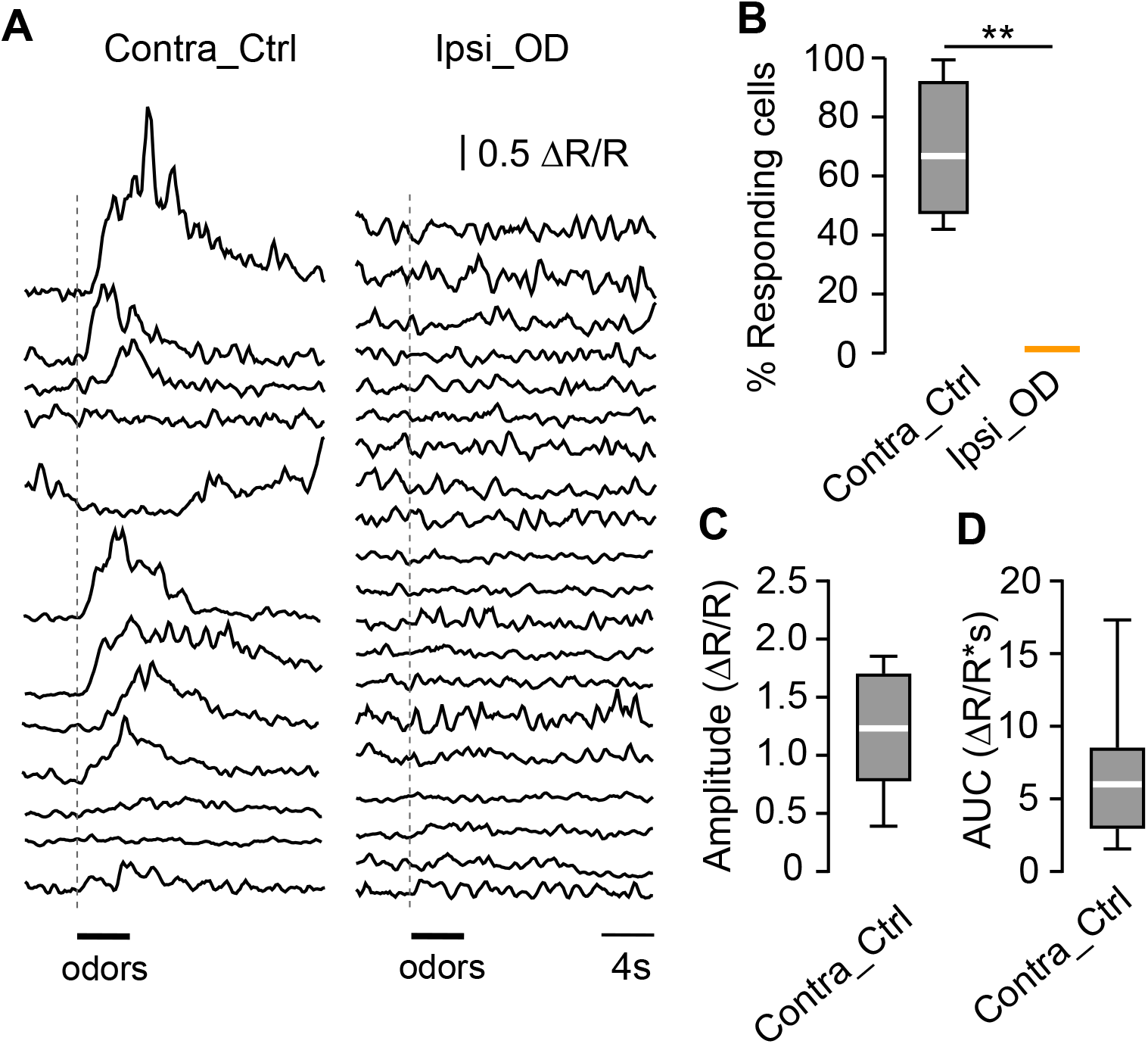
related to Figure1. Nostril occlusion blocked odor-evoked responsiveness of adult-born JGCs. (A) Sample odor-evoked Ca^2+^ transients of adult-born JGCs residing in the contralateral control and ipsilateral odor-deprived hemibulbs. (B-D) Box plots illustrating the fractions of odor responding cells per mouse (B) and (per cell) the amplitude (C) and AUC (D) of odor-evoked Ca^2+^ transients. n=13/4 and 29/4 cells/mice for contralateral control and ipsilateral odor-deprived hemibulbs. Data are shown as median ± IQR. \*\**P*<0.01.

**Figure S5.**
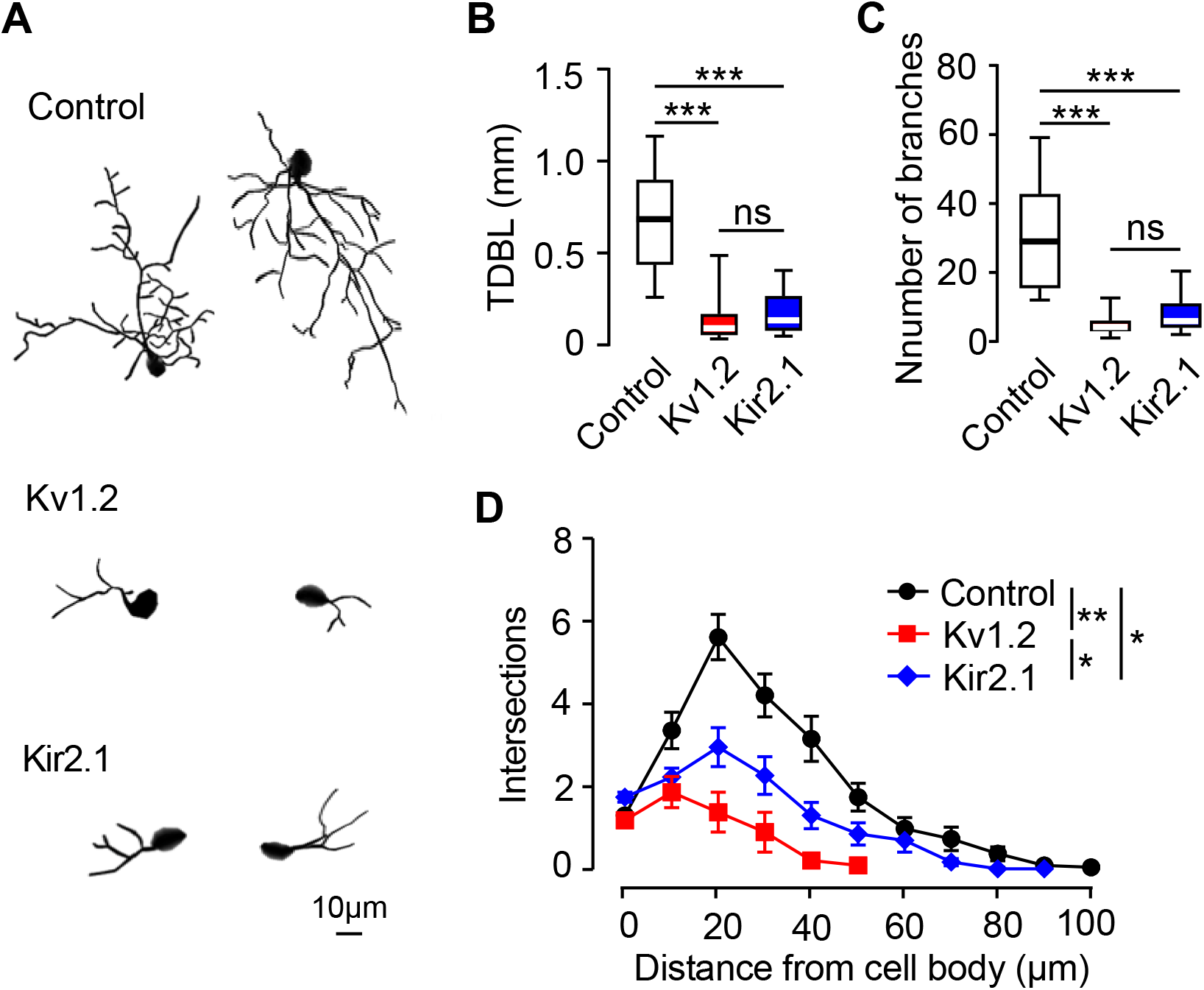
related to Figure 2. Immunocytochemical analyses confirming the retarded morphology of Kv1.2- and Kir2.1-expressing adult-born JGCs. (A) Sample images of adult-born JGCs (20 dpi) belonging to control, Kv1.2 and Kir2.1 groups, reconstructed from *in vitro* images taken in 50 µm thick fixed slices, immunofluorescently labeled against Twitch-2B. (B) Box plots showing (per cell) the TDBL of adult-born JGCs. (C) Box plots showing (per cell) the number of dendritic branches of adult-born JGCs. (B) and (C): n=26/5, 25/5, and 25/5 cells/mice for control, Kv1.2, and Kir2.1, respectively. (D) Sholl analysis illustrating the complexity of adult-born JGC’s dendritic morphology in fixed brain slices at 20 dpi. n=25/5 cells/mice per group. (B) and (C): Data are shown as median ± IQR. (D) Data are shown as mean ± SEM. \**P*<0.05, \*\**P*<0.01, \*\*\**P*<0.001, ns=not significant.

**Figure S6.**
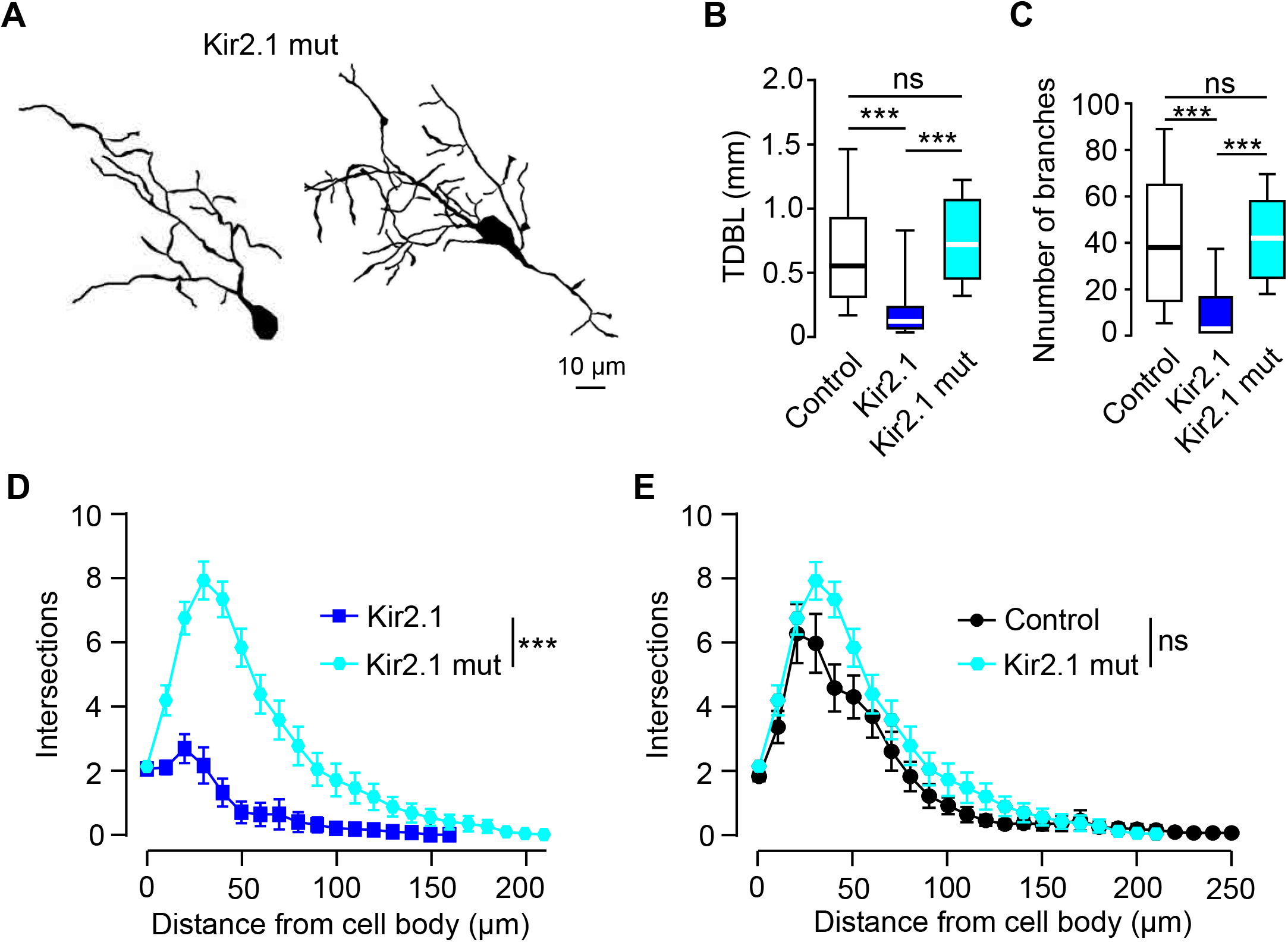
related to Figure 2. Expression of Kir2.1 channel with a dominant-negative loss-of-function mutation (Kir2.1mut) did not impair the morphogenesis of adult-born JGCs. (A) Sample reconstructions of *in vivo* adult-born JGCs (20 dpi) expressing Kir2.1mut. (B) Box plots showing (per cell) the TDBL of adult-born JGCs imaged *in vivo* at 20 dpi in control, Kir2.1, and Kir2.1mut groups. (C) Box plots showing (per cell) the number of dendritic branches of adult-born JGCs imaged *in vivo* at 20 dpi in control, Kir2.1, and Kir2.1mut groups. Control and Kir2.1 datasets are the same as in Figure 2. (D) Sholl analysis of Kir2.1- and Kir2.1mut-expressing adult-born JGCs. (E) Sholl analysis of control and Kir2.1mut-expressing adult-born JGCs. (D) and (E): n=33/7, 46/6, and 35/3 cells/mice for control, Kir2.1, and Kir2.1mut groups. (B) and (C): Data are shown as median ± IQR. (D) and (E): Data are shown as mean ± SEM. \*\*\**P*<0.001, ns=not significant.

**Figure S7.**
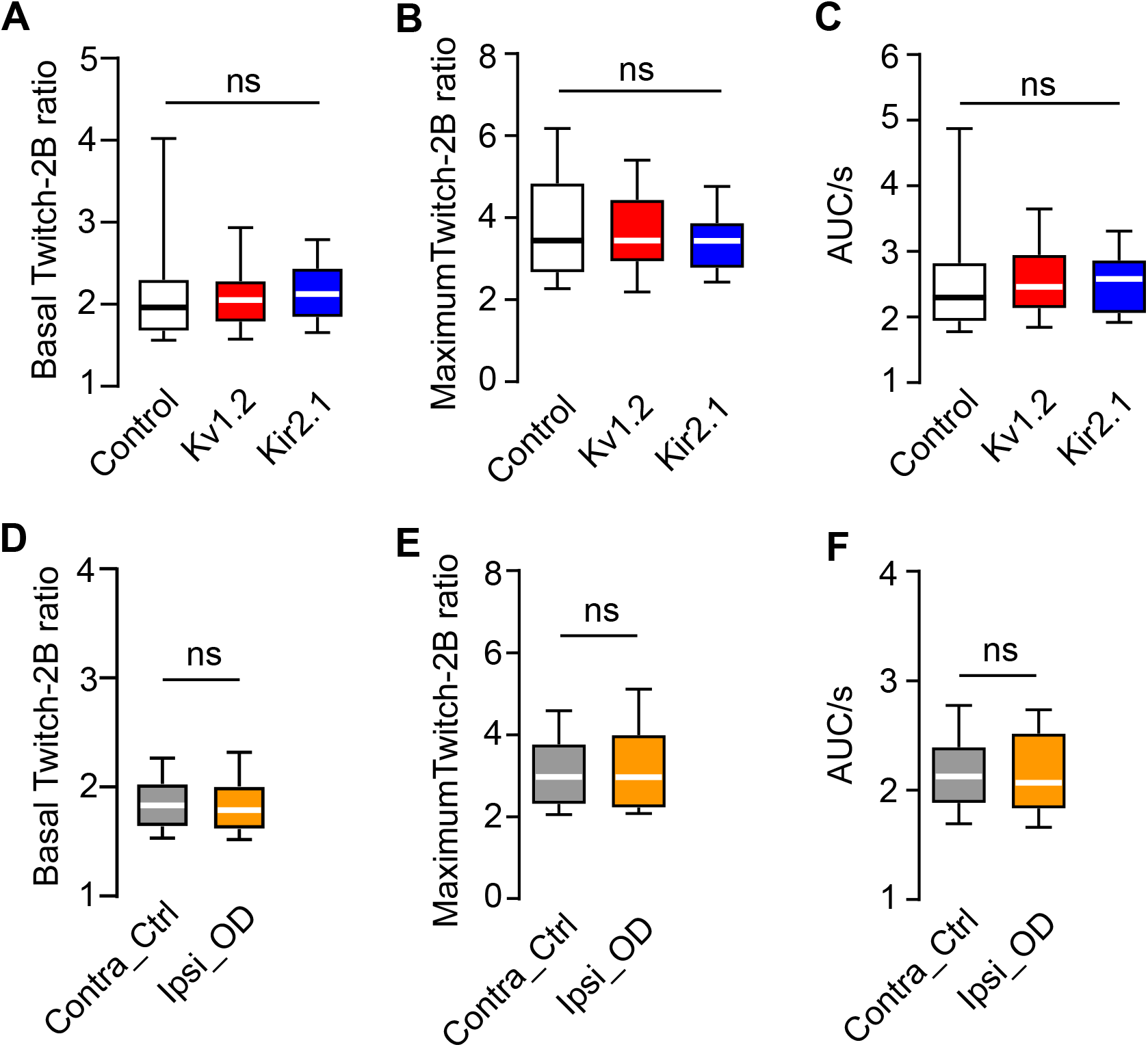
related to Figure5. Neither Kv1.2 or Kir2.1 overexpression nor odor deprivation affected basal and maximum Twitch-2B ratios and AUCs of adult-born JGCs. Box plots showing the median (per cell) basal and maximum Twitch-2B ratios and AUCs of spontaneous Ca^2+^ transients in adult-born JGCs in control, Kv1.2, and Kir2.1 groups (A-C) as well as contralateral control and ipsilateral odor-deprived groups (D-F). n=60/8, 68/7, 66/5, 34/4, and 47/4 cells/mice for control, Kv1.2, Kir2.1, contralateral control, and ipsilateral odor-deprived groups, respectively. Data are shown as median ± IQR. ns=not significant.

**Figure S8.**
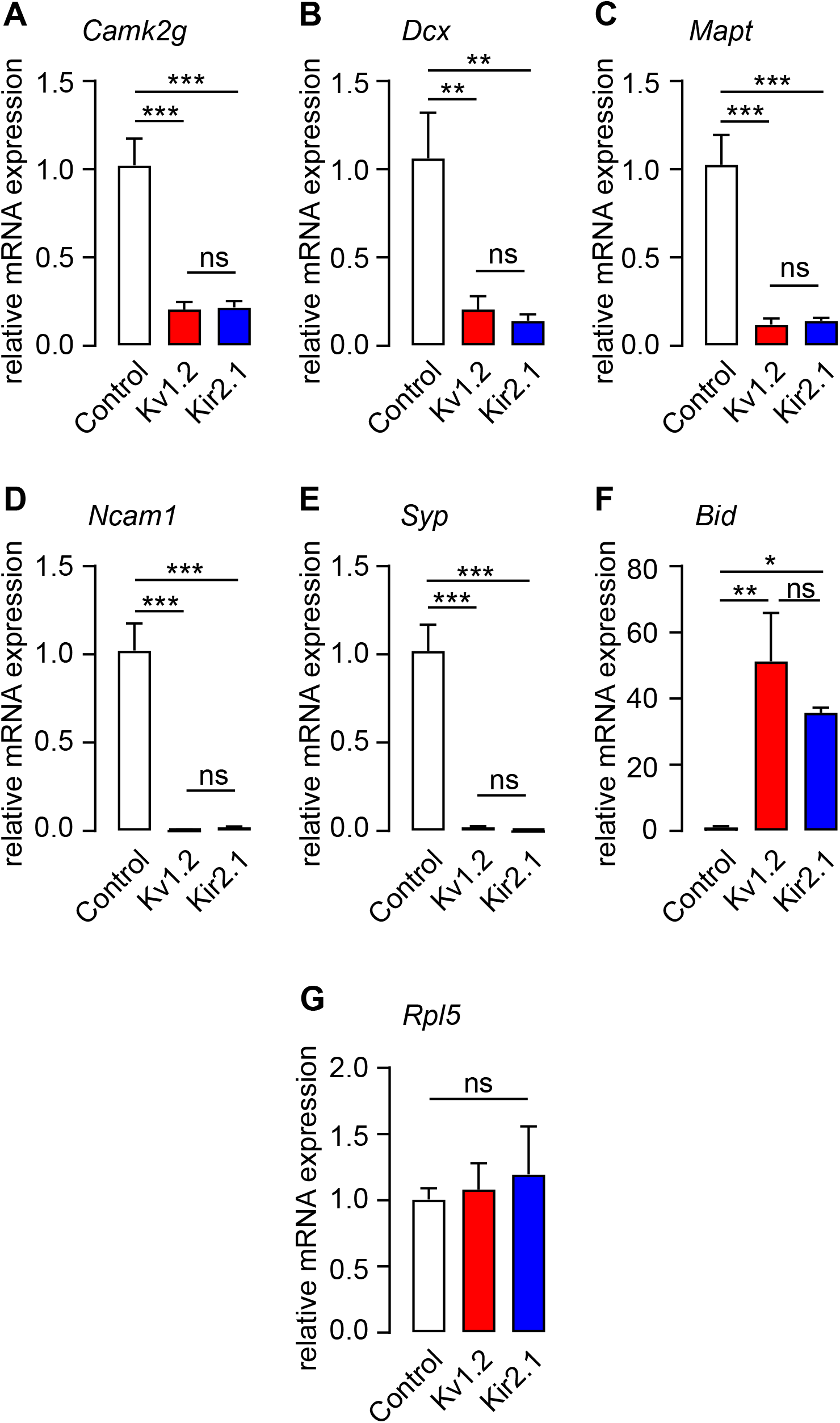
related to Figure 7. Relative expression levels of transcripts involved in pathways related to neuronal development, differentiation, and cell death. Bar graphs showing mean relative expression levels of mRNA encoding for the calcium/calmodulin-dependent protein kinase II gamma (*Camk2g*; A), doublecortin (*Dcx*; B), microtubule-associated protein tau (*Mapt*; C), neural cell adhesion molecule 1 (*Ncam1*; D), synaptophysin (*Syp*; E), BH3 interacting domain death agonist (*Bid*; F) and ribosomal protein L5 (*Rpl5*; G), as determined by qPCR analyses of FACS-sorted adult-born cells (see Methods). Data are shown as mean ± SEM. n = 3 biological replicates. \**P* < 0.05, \*\**P*<0.01, \*\*\**P*<0.001, ns=not significant.

**Figure S9.**
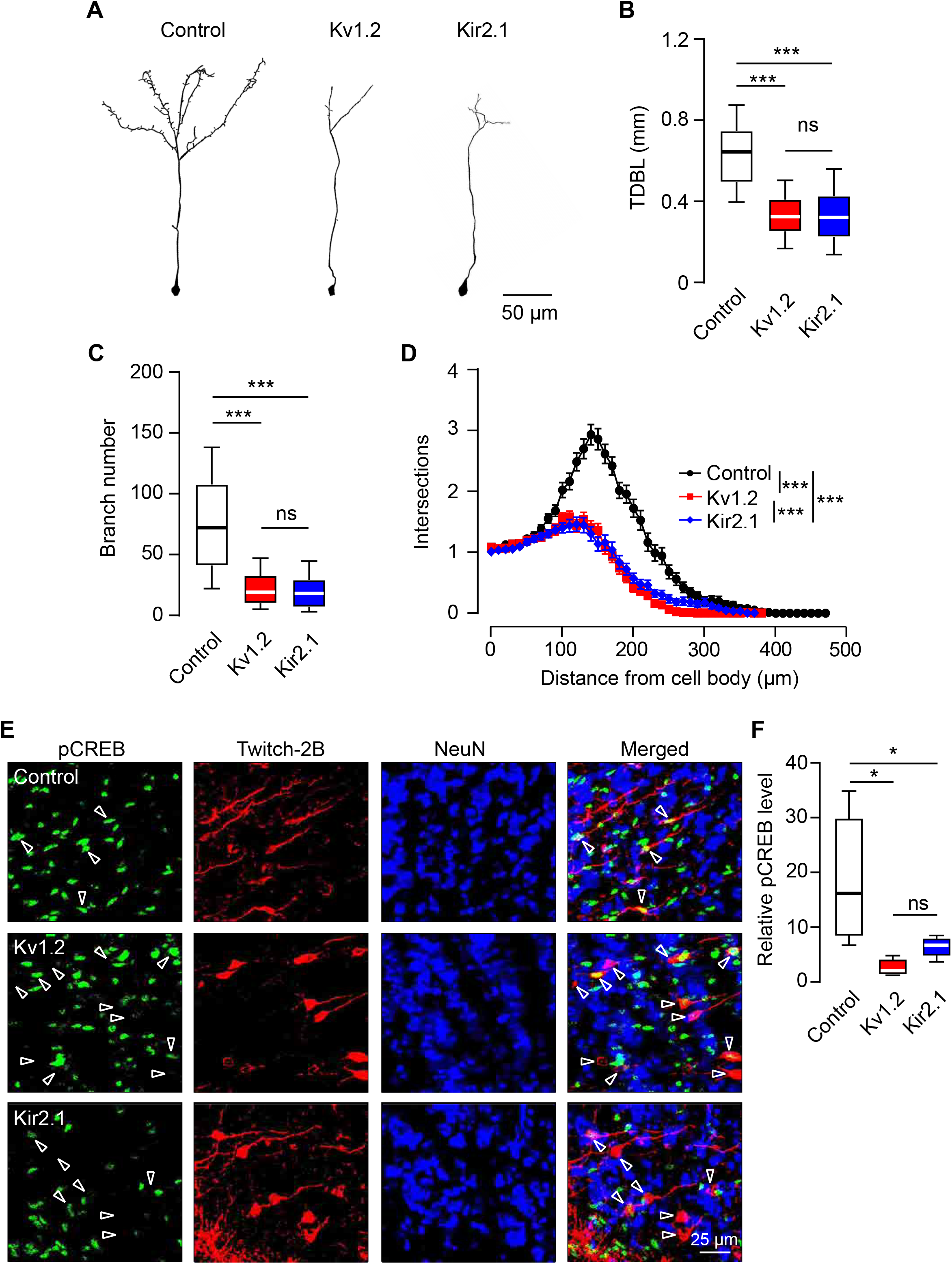
related to Figures 2, 6 and 7. Morphology and relative pCREB levels of adult-born granule cells from control, Kv1.2, and Kir2.1 groups. (A) Sample images of adult-born GCs (20 dpi) belonging to control, Kv1.2 and Kir2.1 groups, reconstructed from *in vitro* images taken in 50 µm thick fixed slices, immunofluorescently labeled against Twitch-2B. (B) Box plots showing (per cell) the TDBL (B) and the number of dendritic branches (C) of adult-born GCs. (D) Sholl analysis illustrating the complexity of adult-born GCs’ dendritic morphology in fixed brain slices at 20 dpi. (E) Sample MIP images (15 µm depth) showing pCREB-, Twitch-2B- and NeuN-positive cells in the granule cell layer of the OB slices from control, Kv1.2 and Kir2.1 groups at 10 dpi. Twitch-2B labels the adult-born GCs whereas NeuN labels the mature neurons. Arrowheads highlight the location of adult-born JGCs. (F) Box plots illustrating the median (per mouse) relative pCREB levels of adult-born GCs in control, Kv1.2, and Kir2.1 groups at 10 dpi. n=119/5, 207/4, and 222/5 cells/mice for control, Kv1.2 and Kir2.1, respectively. (B-D): n=177/7, 149/5, and 133/6 cells/mice for control, Kv1.2 and Kir2.1 respectively. (B) (C) and (F): Data are shown as median ± IQR. (D): Data are shown as mean ± SEM. \**P*<0.05, \*\*\**P*<0.001, ns=not significant.

**Figure S10.**
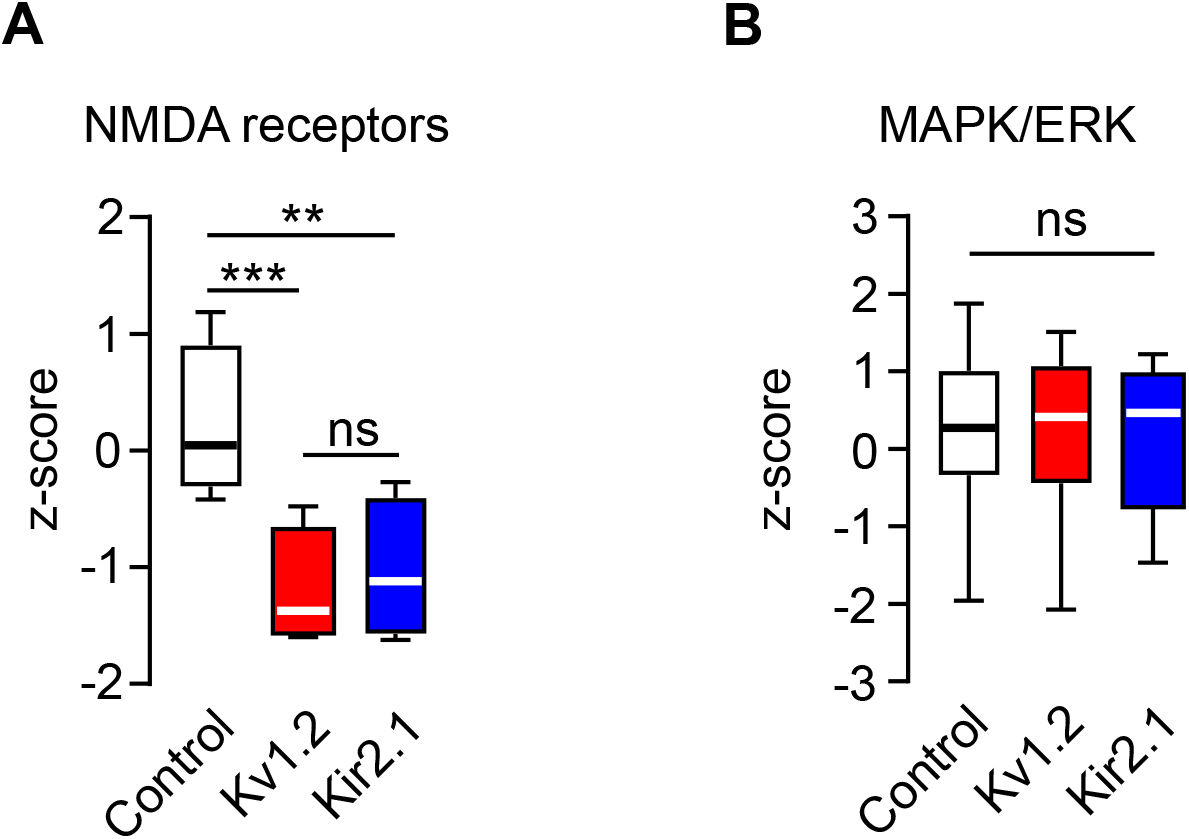
related to Figure 7. Relative expression levels of transcripts encoding for N-methyl-D-aspartate (NMDA) receptors and members of mitogen-activated protein kinase (MAPK) signaling cascade. Box plots showing mean values of z-scores for four isoforms of the NMDA receptor (A) and 18 members of the MAPK/ERK signaling cascade (B). All transcripts included in the box plots are listed in Table S1. n = 2 biological replicates (12 mice in total, 2 mice per replicate per group). Data are shown as median ± IQR. \*\**P*<0.01, \*\*\**P*<0.001, ns=not significant.

**Table S1.**
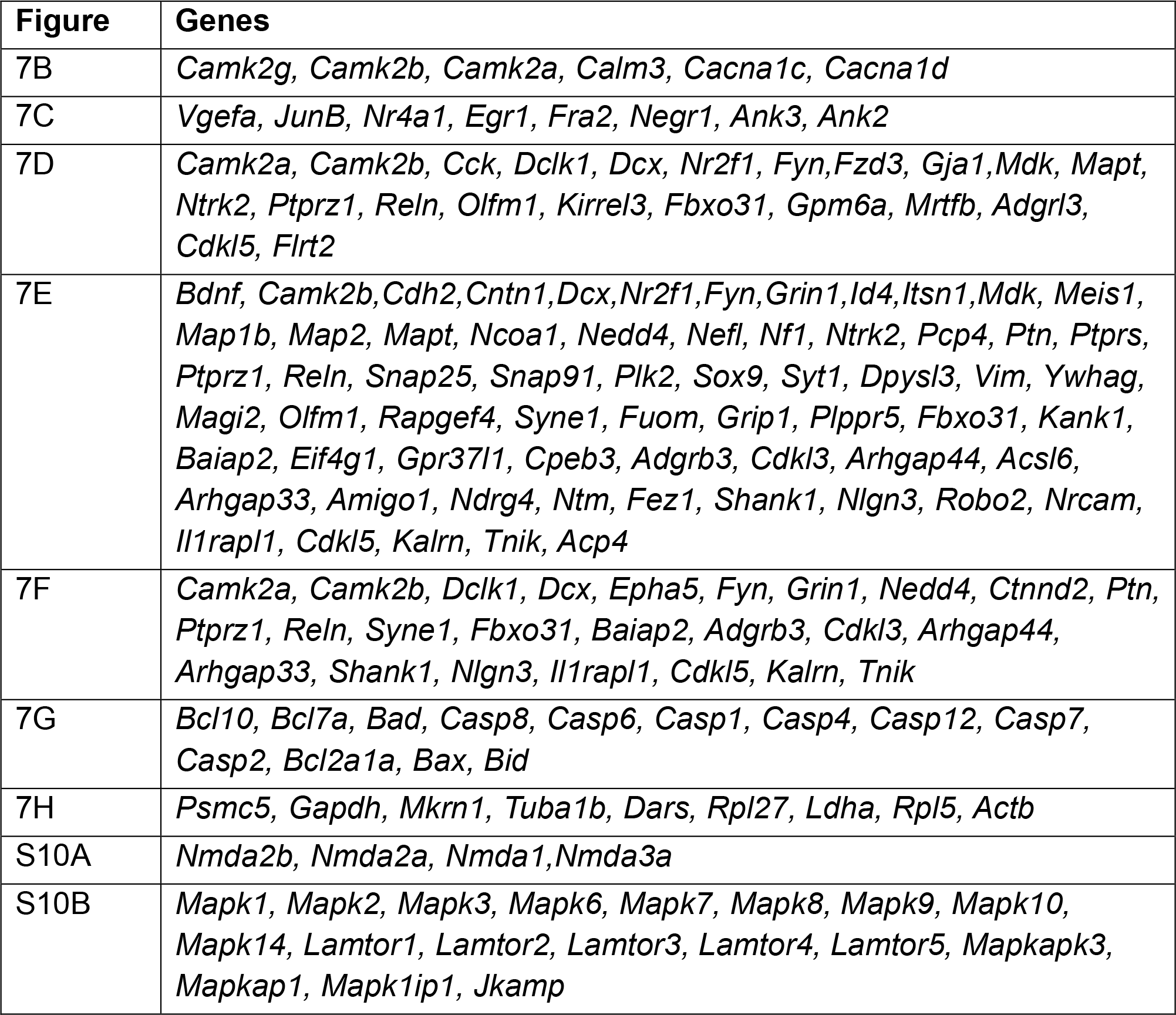
Transcripts included in the box plots shown in Figures 7 and S10.

**Table S2.** related to Figure 7. Differential expression of factors interacting with CREB.

| **Genes** | **Control vs Kv1.2** |  | **Control vs Kir2.1** |  |
| --- | --- | --- | --- | --- |
|  | logFC | P value | logFC | P value |
| *Crtc2* | 4.59 | 0.1818 | 6.34 | 0.1002 |
| *Crtc3* | 0.21 | 0.9484 | 0.61 | 0.8533 |
| *Lmo4* | -1.65 | 0.2554 | -1.04 | 0.4539 |
| *Dream (Kcnip3)* | -0.84 | 0.5959 | -8.61 | 0.0046 |
| *Crest (Ss18l1)* | -6.04 | 0.1098 | 0.56 | 0.8377 |
| *Smarca4* | -1.55 | 0.5495 | -0.91 | 0.7210 |
| *Hdac1* | -1.84 | 0.3210 | 0.29 | 0.8663 |

**Table S3.**
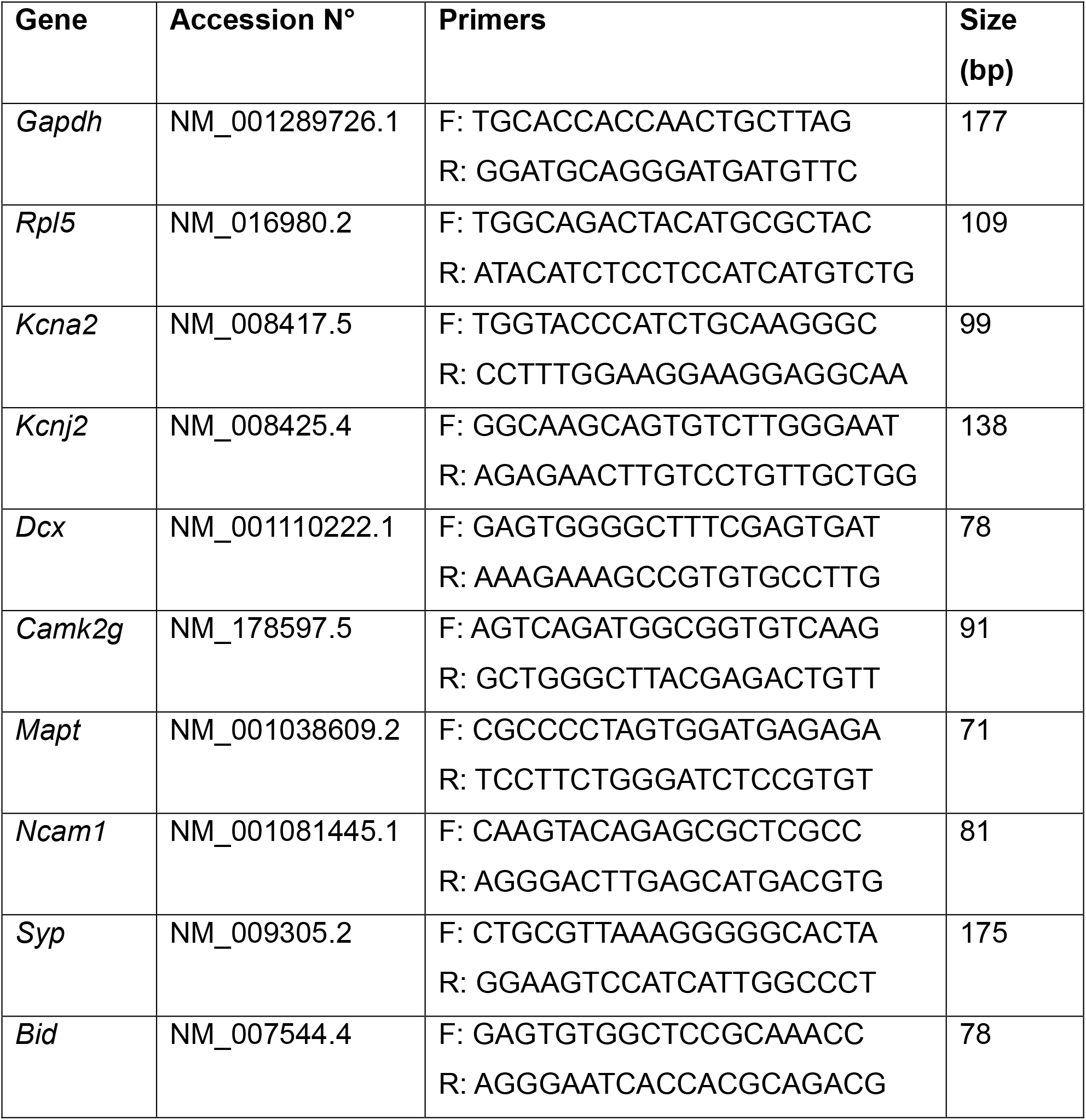
related to Figure S8. Primers used for qPCR. mRNA sequence accession numbers were obtained from the Mouse Genome Database accessed at www.ncbi.nlm.nih.gov/genome.

**Table S4.** Exact results of all statistical tests

| Figure | Statistical test | *Post hoc* comparisons |
| --- | --- | --- |
| 1C | Kruskal-Wallis test: $P=2\times 10^{-4}$ | Dunn's multiple comparison test: control vs. Kv1.2: $P=0.04$, control vs. Kir2.1: $P=2\times 10^{-4}$, Kv1.2 vs. Kir2.1: $P=0.22$. |
| 1D | Kruskal-Wallis test: $P=6\times 10^{-5}$ | Dunn's multiple comparison test: control vs. Kv1.2: $P=2\times 10^{-4}$, control vs. Kir2.1: $P=1.6\times 10^{-3}$, Kv1.2 vs. Kir2.1: $P=0.36$. |
| 1E | Kruskal-Wallis test: $P=2\times 10^{-4}$ | Dunn's multiple comparison test: control vs. Kv1.2: $P=3\times 10^{-4}$, control vs. Kir2.1: $P=6.3\times 10^{-3}$, Kv1.2 vs. Kir2.1: $P=0.67$. |
| 1F | Kruskal-Wallis test: $P=3\times 10^{-4}$ | Dunn's multiple comparison test: control vs. Kv1.2: $P=8.9\times 10^{-3}$, control vs. Kir2.1: $P=3\times 10^{-3}$, Kv1.2 vs. Kir2.1: $P=0.55$. |
| 1G | One-way ANOVA:<br>$F_{2,19}=7.982$, $P=3\times 10^{-3}$ | Tukey's multiple comparison test: control vs. Kv1.2: $P=0.01$, control vs. Kir2.1: $P=6.6\times 10^{-3}$, Kv1.2 vs. Kir2.1: $P=0.59$. |
| 1I-M | Two-sided unpaired $t$ test,<br>$P>0.05$ for all comparisons. | |
| 2B | Kruskal-Wallis test: $P=2\times 10^{-9}$ | Dunn's multiple comparison test: control vs. Kv1.2: $P=3\times 10^{-9}$; control vs. Kir2.1: $P=8\times 10^{-6}$; Kv1.2 vs. Kir2.1: $P=0.21$ |
| 2C | Kruskal-Wallis test: $P=5\times 10^{-10}$ | Dunn's multiple comparison test: control vs. Kv1.2: $P=2\times 10^{-9}$; control vs. Kir2.1: $P=5\times 10^{-7}$; Kv1.2 vs. Kir2.1: $P=0.58$ |
| 2D | Generalized linear mixed model fit by maximum likelihood (Laplace Approximation) | control vs. Kv1.2: $P<0.001$, control vs. Kir2.1: $P<0.001$, Kv1.2 vs. Kir2.1: $P<0.001$ |
| 2F, G | Two-sided unpaired $t$ test,<br>$P>0.05$ for both comparisons. | |
| 2H | Generalized linear mixed model fit by maximum likelihood (Laplace Approximation) | $P=0.98$ |
| 3B | One-way ANOVA:<br>$F_{2,15}=6.747$, $P=8.1\times 10^{-3}$ | Tukey's multiple comparison test: control vs. Kv1.2: $P=0.02$; control vs. Kir2.1: $P=9.6\times 10^{-3}$; Kv1.2 vs. Kir2.1: $P=0.69$. |
| 3D | One-way ANOVA:<br>$F_{2,11}=13.84$, $P=1\times 10^{-3}$ | Tukey's multiple comparison test: control vs. Kv1.2: $P=0.81$; control vs. Kir2.1: $P=1.2\times 10^{-3}$; Kv1.2 vs. Kir2.1: $P=3.3\times 10^{-3}$. |
| 4A | One-way ANOVA:<br>$F_{2,16}=7.061$, $P=0.0063$ | Tukey's multiple comparison test: control vs. Kv1.2: $P=0.54$; control vs. Kir2.1: $P=5.9\times 10^{-3}$; Kv1.2 vs. Kir2.1: $P=0.04$. |
| 4C | Kruskal-Wallis test: $P=2\times 10^{-6}$ | Dunn's multiple comparison test: control vs. Kv1.2: $P=4\times 10^{-5}$; control vs. Kir2.1: $P=6\times 10^{-4}$; Kv1.2 vs. Kir2.1: $P>0.99$. |
| 4D | Kruskal-Wallis test: $P=7\times 10^{-9}$ | Dunn's multiple comparison test: control vs. Kv1.2: $P=3\times 10^{-6}$; control vs. Kir2.1: $P=4\times 10^{-6}$; Kv1.2 vs. Kir2.1: $P=0.39$. |
| 5B | Chi-square test, $P=3\times 10^{-6}$ | |
| 5C | One-way ANOVA:<br>$F_{2,15}=5.468$, $P=0.016$ | Tukey's multiple comparison test: control vs. Kv1.2: $P=0.03$; control vs. Kir2.1: $P=6.2\times 10^{-3}$; Kv1.2 vs. Kir2.1: $P=0.57$. |
| 5E | Chi-square test, $P=0.28$ | |
| 5F | Two-sided unpaired $t$ test,<br>$P=0.75$ | |
| 6B | One-way ANOVA:<br>$F_{4,22}=14.66$, $P=6\times 10^{-6}$ | Tukey's multiple comparison test: control vs. Kv1.2: $P=7\times 10^{-5}$; control vs. Kir2.1: $P=2\times 10^{-4}$; control vs. Contra_Ctrl: $P=0.79$; control vs. Ipsi_OD: $P=0.95$; Kv1.2 vs. Kir2.1: $P=0.99$; Kv1.2 vs. Contra_Ctrl: $P=2.1\times 10^{-3}$; Kv1.2 vs. Ipsi_OD: $P=7\times 10^{-4}$; Kir2.1 vs. Contra_Ctrl: $P=3.8\times 10^{-3}$; Kir2.1 vs. Ipsi_OD: $P=1.4\times 10^{-3}$; Contra_Ctrl vs. Ipsi_OD: $P=0.99$. |
| 6C | One-way ANOVA:<br>$F_{2,12}=27.88$, $P=3\times 10^{-5}$ | Tukey's multiple comparison test: control vs. Kv1.2: $P=1\times 10^{-5}$; control vs. Kir2.1: $P=7\times 10^{-5}$; Kv1.2 vs. Kir2.1: $P=0.95$. |
| 7A | One-way ANOVA:<br>$F_{2,10}=14.12$, $P=2.4\times 10^{-3}$ | Tukey's multiple comparison test: control vs. Kv1.2: $P=2.9\times 10^{-3}$; control vs. Kir2.1: $P=2.2\times 10^{-3}$; Kv1.2 vs. Kir2.1: $P=0.99$. |
| 7B | One-way ANOVA:<br>$F_{2,14}=18.02$, $P=1.3\times 10^{-4}$ | Tukey's multiple comparison test: control vs. Kv1.2: $P=4.3\times 10^{-3}$; control vs. Kir2.1: $P=1.1\times 10^{-4}$; Kv1.2 vs. Kir2.1: $P=0.15$. |
| 7C | One-way ANOVA: $F_{2,42}=121.7$, $P=1\times 10^{-15}$, | Tukey's multiple comparison test: control vs. Kv1.2: $P=1\times 10^{-12}$; control vs. Kir2.1: $P=1\times 10^{-12}$; Kv1.2 vs. Kir2.1: $P=0.56$. |
| 7D | Friedman test, $P=1\times 10^{-15}$ | Dunn's multiple comparison test: control vs. Kv1.2: $P=1\times 10^{-15}$; control vs. Kir2.1: $P=1\times 10^{-15}$; Kv1.2 vs. Kir2.1: $P=0.75$. |
| 7E | One-way ANOVA: $F_{2,48}=99.8$, $P=1\times 10^{-15}$ | Tukey's multiple comparison test: control vs. Kv1.2: $P=1\times 10^{-15}$; control vs. Kir2.1: $P=2\times 10^{-15}$; Kv1.2 vs. Kir2.1: $P=0.84$. |
| 7F | One-way ANOVA: $F_{2,26}=22.02$, $P=2.5\times 10^{-6}$ | Tukey's multiple comparison test: control vs. Kv1.2: $P=1.3\times 10^{-5}$; control vs. Kir2.1: $P=1.4\times 10^{-5}$; Kv1.2 vs. Kir2.1: $P=0.99$. |
| 7G | One-way ANOVA: $F_{2,16}=1.27$, $P=0.31$ | |
| S1C | Kolmogorov-Smirnov test, $P=1\times 10^{-8}$, $D=0.399$. | |
| S1D | Unpaired $t$ test with Welch's correction, $t_5=17.13$, $P=1\times 10^{-5}$. | |
| S1E | One-way ANOVA: $F_{2,6}=696.8$, $P=7.9\times 10^{-8}$ | Tukey's multiple comparison test: control vs. Kv1.2: $P=2.4\times 10^{-7}$; control vs. Kir2.1: $P=0.98$; Kv1.2 vs. Kir2.1: $P=2.5\times 10^{-7}$. |
| S1F | One-way ANOVA: $F_{2,6}=8.16$, $P=0.02$ | Tukey's multiple comparison test: control vs. Kv1.2: $P=0.97$; control vs. Kir2.1: $P=0.03$; Kv1.2 vs. Kir2.1: $P=0.03$. |
| S2A | One-way ANOVA: $F_{2,17}=0.05$, $P=0.95$ | |
| S2B | One-way ANOVA: $F_{2,13}=2.18$, $P=0.15$ | |
| S3A | One-way ANOVA: $F_{2,18}=4.01$, $P=0.04$ | unpaired $t$ test with Welch's correction for multiple comparisons: control vs. Kv1.2: $P=0.17$; control vs. Kir2.1: $P=1.9\times 10^{-3}$; Kv1.2 vs. Kir2.1: $P=0.04$. |
| S3B | Two-sided Mann-Whitney test, $P=0.68$ | |
| S3C | Two-sided Mann-Whitney test, $P=0.82$ | |
| S3D | Two-sided Mann-Whitney test, $P=0.02$ | |
| S4B | Unpaired $t$ test with Welch's correction, $t_3=6.31$, $P=8\times 10^{-3}$ | |
| S5B | Kruskal-Wallis test: $P=5\times 10^{-9}$ | Dunn's multiple comparison test: control vs. Kv1.2: $P=5\times 10^{-9}$; control vs. Kir2.1: $P=4\times 10^{-6}$; Kv1.2 vs. Kir2.1: $P=0.72$ |
| S5C | Kruskal-Wallis test: $P=6\times 10^{-10}$ | Dunn's multiple comparison test: control vs. Kv1.2: $P=1\times 10^{-9}$; control vs. Kir2.1: $P=2\times 10^{-5}$; Kv1.2 vs. Kir2.1: $P=0.27$ |
| S5D | Generalized linear mixed model fit by maximum likelihood (Laplace Approximation) | control vs. Kv1.2: $P=0.004$, control vs. Kir2.1: $P=0.011$, Kv1.2 vs. Kir2.1: $P=0.016$ |
| S6B | Kruskal-Wallis test: $P=1\times 10^{-9}$ | Dunn's multiple comparison test: control vs. Kir2.1: $P=1\times 10^{-5}$; control vs. Kir2.1mut: $P=0.7$; Kir2.1 vs. Kir2.1mut: $P=6\times 10^{-9}$ |
| S6C | Kruskal-Wallis test: $P=7\times 10^{-11}$ | Dunn's multiple comparison test: control vs. Kir2.1: $P=3\times 10^{-7}$; control vs. Kir2.1mut: $P=0.99$; Kir2.1 vs. Kir2.1mut: $P=3\times 10^{-9}$ |
| S6D | Generalized linear mixed model fit by maximum likelihood (Laplace Approximation) | $P<0.001$ |
| S6E | Generalized linear mixed model fit by maximum likelihood (Laplace Approximation) | $P=0.28$ |
| 7A-C | Kruskal-Wallis test, $P>0.05$ for all tests | |
| S7D-F | Mann-Whitney test, $P>0.05$ for all tests | |
| S8A | One-way ANOVA: $F_{2,6}=25.21$, $P=0.0012$ | Fisher's multiple comparison test: control vs. Kv1.2: $P=8.2\times 10^{-4}$; control vs. Kir2.1: $P=8.8\times 10^{-4}$; Kv1.2 vs. Kir2.1: $P=0.94$. |
| S8B | One-way ANOVA: $F_{2,6}=10.77$, $P=0.01$ | Fisher's multiple comparison test: control vs. Kv1.2: $P=8.3\times 10^{-3}$; control vs. Kir2.1: $P=6\times 10^{-3}$; Kv1.2 vs. Kir2.1: $P=0.78$. |
| S8C | One-way ANOVA: $F_{2,6}=27.17$, $P=9.8\times 10^{-4}$ | Fisher's multiple comparison test: control vs. Kv1.2: $P=6.5\times 10^{-4}$; control vs. Kir2.1: $P=7.4\times 10^{-4}$; Kv1.2 vs. Kir2.1: $P=0.88$. |
| S8D | One-way ANOVA: $F_{2,6}=43.04$, $P=2.7\times 10^{-4}$ | Fisher's multiple comparison test: control vs. Kv1.2: $P=1.9\times 10^{-4}$; control vs. Kir2.1: $P=2.1\times 10^{-4}$; Kv1.2 vs. Kir2.1: $P=0.91$. |
| S8E | One-way ANOVA: $F_{2,6}=46.98$, $P=2.2\times 10^{-4}$ | Fisher's multiple comparison test: control vs. Kv1.2: $P=1.6\times 10^{-4}$; control vs. Kir2.1: $P=1.5\times 10^{-4}$; Kv1.2 vs. Kir2.1: $P=0.89$. |
| S8F | One-way ANOVA: $F_{2,6}=9.19$, $P=0.015$ | Fisher's multiple comparison test: control vs. Kv1.2: $P=6\times 10^{-3}$; control vs. Kir2.1: $P=0.03$; Kv1.2 vs. Kir2.1: $P=0.24$. |
| S8G | One-way ANOVA: $F_{2,6}=0.15$, $P=0.86$ | |
| S9B | Kruskal-Wallis test: $P=1\times 10^{-14}$ | Dunn's multiple comparison test: control vs. Kv1.2: $P=1\times 10^{-14}$; control vs. Kir2.1: $P=1\times 10^{-14}$; Kv1.2 vs. Kir2.1: $P=0.99$ |
| S9C | Kruskal-Wallis test: $P=1\times 10^{-14}$ | Dunn's multiple comparison test: control vs. Kv1.2: $P=1\times 10^{-14}$; control vs. Kir2.1: $P=1\times 10^{-14}$; Kv1.2 vs. Kir2.1: $P=0.99$ |
| S9D | Generalized linear mixed model fit by maximum likelihood (Laplace Approximation) | control vs. Kv1.2: $P<0.001$, control vs. Kir2.1: $P<0.001$, Kv1.2 vs. Kir2.1: $P<0.001$ |
| S9F | One-way ANOVA: $F_{2,11}=6.524$, $P=0.013$ | Tukey's multiple comparison test: control vs. Kv1.2: $P=0.02$; control vs. Kir2.1: $P=0.049$; Kv1.2 vs. Kir2.1: $P=0.69$. |
| S10A | One-way ANOVA: $F_{2,6}=31.74$, $P=6\times 10^{-4}$ | Tukey's multiple comparison test: control vs. Kv1.2: $P=8\times 10^{-4}$; control vs. Kir2.1: $P=1.6\times 10^{-3}$; Kv1.2 vs. Kir2.1: $P=0.67$. |
| S10B | One-way ANOVA: $F_{2,34}=0.18$, $P=0.83$ | |

## Materials and Methods

### Mouse models

All experimental procedures were performed in accordance with Institutional Animal Welfare Guidelines and were approved by the state government of Baden-Württemberg, Germany. Three-to four-month-old C57BL/6 mice of either sex were used in this study. Animals were kept in pathogen-free conditions at 22 °C, 60% air humidity, 12-hours light-dark-cycle with ad libitum access to food and water. Females stayed in groups of 3-5 mice, males were kept individually. Mice of similar age were assigned randomly to control and test groups. Littermates were evenly distributed among experimental groups and each experimental group contained mice from different litters.

### Implantation of a cranial window

A chronic cranial window was implanted over the mouse OB as described previously(19, 35). Mice were anesthetized by an intraperitoneal (i.p.) injection of ketamine/xylazine (80/4 μg/g of body weight (BW)). Anesthetic depth was monitored by toe pinches throughout the surgery and additional ketamine/xylazine (40/2 μg/g of BW) was injected when necessary. Dexamethasone (2 μg/g of BW) was administered intramuscularly before the surgery. Local anesthetic lidocaine (2%) was applied subcutaneously over the OB 5-10 minutes before removing the scalp. The ointment was used to prevent dehydration of the mouse’s eyes. A circular cranial opening (3 mm in diameter) was made by repeated drilling over the two OB hemispheres. Small pieces of bone were removed with a sharp blade and tweezers. Extreme care was taken, not to damage any blood vessels on the surface of the OB. The opening was rinsed with a standard extracellular solution (composition in mM: 125 NaCl, 4.5 KCl, 26 NaHCO_3_, 1.25 NaH_2_PO_4_, 2 CaCl_2_, 1 MgCl_2,_ and 20 glucose; pH 7.4, bubbled continuously with 95% O_2_ and 5% CO_2_) and covered with a glass coverslip (Ø 3 mm, Warner Instruments, Hamden, CT, USA). The gap between the edge of the coverslip and the skull was filled with cyanoacrylate glue and then strengthened by dental cement. During the surgery and until full recovery from anesthesia the mouse was kept on a heated plate. Postoperative care included an analgesic dose of carprofen (5 μg/g of BW) for 3 days subcutaneously and the antibiotic Enrofloxacin (1:100 v/v) in drinking water for consecutive 10 days. Mice were allowed to recover for at least 3-4 weeks and were subsequently examined for window clarity. Mice were singly housed after cranial window implantation on a 12 h light/dark cycle with food and water available *ad libitum*.

### Construction of viral vectors and production of viruses

All lentiviral vectors were based on the FUGW backbone(72). The eGFP in the original FUGW plasmid was replaced by a Ca^2+^ indicator Twitch-2B at BamHI and EcoRI restriction sites to generate FUW-Twitch-2B (73). Kv1.2wt-T2A and Kir2.1wt-T2A fragments were magnified by PCR (Phusion High-Fidelity PCR Kit, NEB) from pcDNA3-Kv1.2wt (human cDNA template) and Mgi-Kir2.1wt (mouse cDNA template, from Carlos Lois laboratory, Caltech) plasmids and were inserted into FUW-Twitch-2B between XbaI and BamHI restriction enzyme sites. To construct Lenti-Twitch-2B-T2A-H2B-mCherry, H2B-mCherry was inserted after Twitch-2B and T2A to enable nuclear-located expression of mCherry. The sequence of T2A used in this study is 5’-tgggccaggattctcctcgacgtcaccgcatgttagcagacttcctctgccctctccactgcctaccgg-3’.

The second generation of virus packaging system was used in this study. Briefly, HEK-293T cells were transiently transfected with plasmids encoding target genes (Lenti-Twitch-2B-T2A-H2B-mCherry, Lenti-Kv1.2-T2A-Twitch-2B, or Lenti-Kir2.1-T2A-Twitch-2B), plus viral packaging helper plasmids pMD2.G (plasmid #12259, Addgene) and psPAX2 (plasmid #12260, Addgene) as described previously (35). The Lipofectamine 3000 reagent (Invitrogen) was used for transfection. HEK-293T and 1F8 cells were cultured in Dulbecco’s Modified Eagle’s Medium (DMEM) with 10% heat-inactivated fetal bovine serum and 2 mM L-glutamine at 37°C and 5% CO_2_ in air atmosphere. Cell lines were not authenticated. Mycoplasma contamination was controlled by regular PCR tests. To block the K^+^ channel overexpression-induced reduction of the lentivirus-producing capacity of HEK-293T cells (74), K^+^ channel blocker Ba^2+^ (0.3 mM) was added to the cell culture medium. 48-72 hours after transfection, cell culture supernatant containing viral particles was collected and concentrated by centrifugation at 135,000 g at 4°C for 2 hours. Concentrated supernatants were resuspended with PBS and titrated in HEK-293T cells. Titers of about 8×10^9^ virus particles per ml concentrated supernatant were used for the following experiments. Retroviral vector Mgi-Kir2.1mut encoding 3 dominant negative site mutations (G144A, Y145A, G146A; Carlos Lois laboratory, Caltech), was packaged in 1F8 cells derived from 293GPG cell line. The vector was derived from a Moloney leukemia virus with an internal promoter from the Rous sarcoma virus. 48 hours after transfection, cell culture supernatant containing the retroviral particles was harvested and concentrated by centrifugation at 135,000 g at 4°C for 6 hours. After concentration, the pellet was resuspended with as little volume of ice-chilled PBS as possible. Titers above 10^8^ particles per ml supernatant were used for *in vivo* injections.

### Virus injection into the RMS

Animals with implanted cranial windows were anesthetized with ketamine/xylazine (80/4 μg/g of BW) and fixed in a stereotaxic frame. The ointment was used to prevent dehydration of the mouse’s eyes and 2% lidocaine was applied subcutaneously on top of the injection sites. Viruses were stereotactically injected into the RMS at the following coordinates: anterior-posterior +3.0 mm, medial-lateral ±0.83, and dorsal-ventral −2.95 ±0.05 mm from the pial surface. Approximately, 0.8-1 μl of the virus-containing solution was injected into the RMS of each hemisphere. Thereafter, a metal bar, required for head fixation during the subsequent imaging sessions, was fixed to the caudal part of the skull with dental cement. The other exposed parts of the skull were also covered with dental cement. The mice were returned to the home cage and carprofen (5 μg/g of BW) was injected subcutaneously for 3 subsequent days.

### *In vivo* two-photon imaging

Mice with implanted cranial windows were anesthetized with either isoflurane or an MMF (medetomidine 0.5 μg/g BW, midazolam 5.0 μg/g BW, fentanyl 0.05 μg/g BW) anesthesia and placed on a heating plate. Breathing rate and body temperature were monitored continuously using the animal monitoring system (AD Instruments, Sydney, Australia). The head of the mouse was fixed with the metal bar to the *X*-*Y* table, ensuring consistent positioning through imaging sessions. *In vivo* two-photon imaging was performed using an Olympus FV1000 system (Olympus, Tokyo, Japan) with a MaiTai Deep See Laser (Spectra-Physics, Mountain View, CA, USA) and a Zeiss 20x water-immersion objective lens (NA 1.0, Carl Zeiss, Jena, Germany). Unless otherwise indicated, cells were imaged using the 890 nm excitation wavelength.

#### Imaging migration of adult-born JGCs

Mice were anesthetized with isoflurane (2% for induction, 0.8-1.0% for maintenance) and transferred into the imaging setup. The body temperature was kept at ∼37°C. Breathing rate was monitored during the whole imaging session and maintained at 110-140 breaths per minute by slightly adjusting the isoflurane concentration in O_2_. To measure the migration speed, the positions of adult-born neurons were monitored every 15 minutes for 4 hours at 8 and 14 dpi according to the previously established protocol (35). To create landmarks for single-cell tracking, we labeled blood vessels via i.p. injection of sulforhodamine B (0.1 ml/20g BW, 1 mM in PBS, Sigma-Aldrich, St. Louis, USA). In addition, the same FOVs were re-imaged with the same imaging settings at 11, 25, and 45 dpi.

#### Recording spontaneous and odor-evoked Ca^2+^ transients of adult-born JGCs

The spontaneous Ca^2+^ transients were recorded in awake mice. Prior to imaging sessions, the mice were trained for head fixation for 10-12 days, as described in (13). Spontaneous Ca^2+^ transients of adult-born JGCs were recorded at 12 dpi continuously for 2 minutes with a frame rate of 7-10 Hz. Twitch-2B was excited at 890 nm and the emitted light was split into 2 channels by a 515 nm dichroic mirror. The emission light of mCerulean3 was filtered with a 475/64 nm band-pass filter and the emission light of cpVenus^CD^ was filtered with a 500 nm long-pass filter.

The odor-evoked responsiveness of adult-born JGCs was measured at 20 dpi. Mice were anesthetized using MMF anesthesia, the temperature was kept at ∼37°C and the breathing rate was ∼140 breaths per minute during the whole imaging session. Odors were applied through a custom-built flow-dilution olfactometer, positioned in front of the mouse’s snout as described previously (75). Odor mixture containing 2-hexanone, isoamyl acetate, and ethyl tiglate (purchased from Sigma-Aldrich, 0.6% of saturated vapor each) were applied as a 4-second-long pulse with an inter-pulse interval of at least 2 minutes. The odor delivery was not timed relative to respiration. Each cell was stimulated at least twice, as described in (75).

### Odor deprivation (OD) and two-photon imaging of odor-deprived mice

Unilateral naris closure was performed at 5 dpi. Nose plugs were constructed of 2 mm polyethylene tube (0.58 mm inner diameter, 0.96 mm outer diameter, Portex, UK) and suture thread (size 3-0, Ethicon, Germany) as described previously(35). Mice were anesthetized using MMF anesthesia and the plug was accurately inserted into the nostril. Thereafter, mice received an antidote containing flumazenil (0.5 mg/kg BW, Fresenius, Germany) and atipamezole (2.5 mg/kg BW, Alfavet, Germany) and returned to their home cages. After naris occlusion, physiological conditions of experimental animals (e.g., breathing, body weight, and stability of nose plug) were monitored carefully every day. Two-photon imaging of odor-deprived mice was conducted under MMF anesthesia, as described above.

### Immunocytochemistry

Mice were transcardially perfused with 4% paraformaldehyde (PFA) in PBS. The brains were removed and fixed in 4% PFA for 24 hours at 4°C, and then cryoprotected in 25% sucrose in PBS overnight at 4°C. Next, the brains were embedded in Tissue Tek (Sakura, Zoeterwoude, Netherlands) and frozen at −80°C. The immunostaining was performed on free-floating sagittal cryoslices (thickness 30-50 μm) at room temperature. The sections were incubated in a blocking buffer containing 5% normal donkey serum (Jackson Immuno Research, Dianova) and 0.1% Triton-X 100 (Sigma, USA) in PBS for 1 hour to prevent nonspecific background staining. After blocking, the sections were incubated with the primary antibodies diluted in the blocking buffer. Following primary antibodies were used: goat polyclonal antibody against GFP (Rockland 600-101-215, 1:1,000), mouse monoclonal antibody against Kv1.2 (NeuroMab 75-008, 1:200), rabbit monoclonal antibody against pCREB (Cell Signaling 9198S, 1:400), mouse monoclonal antibody against NeuN (Millipore MAB377, 1:1,000). After overnight incubation with primary antibodies at 4°C, the sections were rinsed in PBS three times for 10 minutes each and incubated with secondary antibodies (2% BSA and 1% Triton-X 100 in PBS) for 2 hours in the dark at room temperature. The secondary antibodies were as follows: donkey-anti-mouse or anti-rabbit IgG-conjugated Alexa Fluor 488 (A21202 or A21206, 1:1,000), donkey-anti-goat IgG-conjugated Alexa Fluor 594 (A11058, 1:1,000), donkey-anti-mouse IgG conjugated Alexa Fluor 680 (A10038, 1:1,000), all purchased from Invitrogen (Grand Island, NY). Afterward, the sections were washed three times in PBS for 10 minutes, transferred to Superfrost Plus charged glass slides (Langenbrink, Emmendingen, Germany), and mounted in Vectashield (Vector Laboratories, USA) or ProLong Gold (Invitrogen) mounting medium. Immunostained slices were imaged using an Olympus Fluoview 300 laser scanning microscope (Olympus, Tokyo, Japan) coupled with a MaiTai mode-locked laser (Spectra Physics, Mountain View, CA, USA). Alexa Fluor 488 and 594 were excited at 800 nm and the emitted light was split by a 570 nm dichroic mirror and filtered with a 536/40 nm band-pass filter as well as a 570 nm long-pass filter. Alexa Fluor 680 was also excited at 800 nm and the signal was collected in the long-pass channel of a 670 nm dichroic mirror.

### RNA sequencing

Male 3-month-old mice were bilaterally injected into the RMS with either Lenti-Twitch-2B-T2A-H2B-mCherry, Lenti-Kv1.2-T2A-Twitch-2B, or Lenti-Kir2.1-T2A-Twitch-2B viruses and sacrificed at 9 dpi. Mice were deeply anesthetized by ketamine/xylazine (100/10 μg/g of BW) and transcardially perfused with 20 ml ice-cold PBS to empty blood vessels from the blood. Then mice were decapitated, and the olfactory bulbs were quickly dissected and transferred into an ice-cold 35 mm dish containing 0.5 ml dissection medium (Hank’s balanced salt solution containing 15 mM HEPES, 25 mM glucose, 0.4 mg/ml DNase I, and 80 U/ml RNAse inhibitor). Olfactory bulbs from two animals were pooled to assure isolation of at least 3000 adult-born cells. Minced tissue was very gently homogenized in an ice-cold dissection medium and passed through a 70 µm cell strainer. The filtrate was spun for 10 min at 250 g in a refrigerated centrifuge at 4°C. Pellet was resuspended in 37% Percoll and centrifuged for 30 min at 800 g and 4°C. First, the upper myelin layer and then supernatant were removed, the pellet was resuspended in sorting buffer (dissection medium without DNase I) and centrifuged once again for 10 min at 800 g and 4°C. Finally, isolated cells were resuspended in sorting buffer and kept on ice until sorting. Adult-born cells expressing Twitch-2B were separated by fluorescence-activated cell sorting on a Sony SH800Z sorter (Sony Biotechnology Inc, Surrey, UK) immediately after staining samples with propidium iodide (PI). Single cells were selected based on FSC-W/FSC-H gating, dead cells were excluded based on the PI-signals and transduced cells were identified by their Twitch-2B signal (using a 450/50 bandpass filter). Single Twitch-2B-expressing cells were sorted directly into Eppendorf tubes containing 10 µl sorting buffer, immediately frozen in liquid nitrogen, and kept at −80°C until libraries for RNA-sequencing analysis were prepared.

The synthesis of the cDNA was performed using the SMART-Seq v4 Ultra Low Input RNA Kit (Takara Bio). 3,000 to 4,500 frozen sorted cells were lysed using a concentration of 100 cells per µl of lysis buffer. Lysis was performed by resuspending the cells by pipetting and incubating for 5 minutes at room temperature. First-strand cDNA synthesis was performed using 5 µl of cDNA for 90 min at 42°C. Amplification of the full-length double-strand cDNA was monitored by qPCR and was stopped at 17 PCR cycles during the linear amplification phase. The resulting cDNA presented a fragment size distribution of 1,500 up to 5,000 bp on the Bioanalyzer High Sensitivity DNA Kit (Agilent) and a concentration above 300 pg/µl measured with Qubit dsDNA HS fluorometric quantification (ThermoFisher Scientific). Next-generation sequencing (NGS) libraries were prepared using 150 pg of cDNA input in the Nextera XT DNA Library Preparation Kit (Illumina), followed by 12 cycles of PCR. Final libraries had a mean fragment size of 370 bp on the Bioanalyzer, a concentration > 5 ng/µl, and a molarity of > 30 nmol/l measured with Qubit. Libraries were sequenced as single reads (75 bp read length) on a NextSeq500 (Illumina) with a depth of > 20 million reads. Library preparation and sequencing procedures were performed by the same individual and we chose a design, aimed to minimize technical batch effects.

### Quantitative real-time PCR

Reverse transcribed RNA from sorted cells was generated by SMART-Seq v4 Ultra Low Input RNA Kit (Takara Bio). Real-time PCR was performed using a Fast SYBR green PCR Master Mix (Applied Biosystems) and the QuantStudio™ 5 equipment (Applied Biosystems, Germany). Specific primers (SI Appendix, Table S3) were designed using the NCBI tool primer BLAST selecting those sequences that span an exon-exon junction to avoid amplification of any contaminating genomic DNA. Three biological samples were determined in triplicate. Single product amplification was confirmed by running the melting curves. Input quantities were normalized to those for glyceraldehyde 3-phosphate dehydrogenase (*Gapdh*) and the relative mRNA expression was estimated by the efficiency corrected method (76).

### Analyses

#### Calculation of the migration speed of adult-born JGCs

The migration speed of adult-born JGCs was analyzed as described previously (35). The 3D image stacks acquired during the consecutive imaging sessions (containing both adult-born JGCs and blood vessels) were aligned offline using the blood vessel pattern as an anatomical landmark. Each cell received an identification number, and its position (the center of the cells’ soma) was identified in each of 17 stacks (0-4 hours, 15 min inter-stack interval), enabling the reconstruction of the cell trajectory. Next, the X-Y coordinates of each cells’ position were read out using the Fluoview 3.0 Viewer (Olympus). For Z-axis coordinates, the depth of the cells was the relative depth from the dura. With 3D coordinates for each position point, migration distance (D) between the two points in space was calculated according to the following formula:

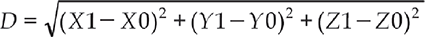

where (X1, Y1, Z1) and (X0, Y0, Z0) were coordinates of the cells’ position in the current and the immediately preceding stacks, respectively. Migration speed was defined as the translocation of the cells’ soma between the two consecutive time points divided by the respective time interval (µm per 15 minutes). Because the step size of the acquired stacks was 2 μm, a cell was considered moving if the translocation of the cells’ soma between the two consecutive time points was more than 4 μm.

#### Analyses of spontaneous Ca^2+^ transients

Data analysis was performed offline with ImageJ and a custom-made routine in Matlab (R2016b, The MathWorks, United States). Circular regions of interest (ROIs) were manually drawn within the soma of each cell. A fluorescence trace for each cell was obtained by averaging all pixels within the ROI. The background signal was obtained from the ROI of the comparable size devoid of fluorescent processes and located near the cell of interest. The image stack was separated into 2 substacks for mCerulean3 and cpVenus^CD^ channels, respectively. The fluorescence traces were calculated separately for both substacks and filtered using a lowpass Butterworth infinite impulse response filter with a cut-off frequency of 0.6 Hz. The Twitch-2B ratio signal was calculated using the formula:

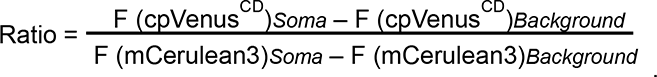

Thereafter, the traces were imported into Matlab to analyze the following parameters: basal and maximum Twitch-2B ratio and the area under the curve normalized to the total recording time (AUC/sec, (37)). The basal ratio was the mean of the lowest 10% data points in the histogram of each individual trace. The value for the maximum ratio was calculated as follows: the filtered traces were processed by a sliding average algorithm with a window size of 1.5 seconds to determine the maximum ratio (maximum average value). Parameters of Ca^2+^ fluctuations were also analyzed in Matlab. The difference between oscillation patterns has been used to compare experimental and control groups. The fluctuations in Ca^2+^ signals were detected using the mid-reference level crossing approach (function midcross in Matlab) and the numbers of crossing points, passing through the mid-reference level, per trace were counted. The Gaussian mixture model(43) was used to explore and detect the naturally existing clusters among the counted numbers of crossing points. The correct number of clusters has been estimated using Bayesian information criterion (77). Assigned cluster labels (i.e., with and without fluctuations in Ca^2+^ signals) have been used to calculate the fraction of each cluster in a given experimental group. Subsequently, these fractions were compared statistically using ANOVA.

#### Analyses of odor-evoked responsiveness of adult-born JGCs

The odor-evoked Ca^2+^ transients of individual neurons were detected with a custom-written Igor Pro routine (WaveMetrics Inc., OR 97035 USA). First, background fluorescence, measured in the neuropil surrounding the adult-born JGCs, was subtracted as described above and the Twitch-2B ratio signals were calculated and expressed as relative Twitch-2B ratio changes (ΔR/R). For automatic detection of responding cells, all ΔR/R traces were smoothed with a binomial filter (time window 0.3 s). Each smoothed trace was subtracted from the original ΔR/R trace, resulting in the “baseline noise” trace. ΔR/R transients were automatically detected with a template-matching algorithm, taking into account their sharp rise. A ΔR/R change was recognized as an odor-evoked Ca^2+^ transient if its amplitude was three times larger than the standard deviation of the corresponding baseline noise.

#### Sholl analyses

The dendritic morphology of adult-born JGCs was analyzed both *in vivo* and *in situ*. The three-dimensional stacks were imported into Neuromantic software (https://www.reading.ac.uk/neuromantic/body_index.php) for 3D reconstruction. Neuronal morphology was manually traced to obtain accurate reconstructions. Digitally reconstructed neurons were imported into Image J and analyzed with the Simple Neurite Tracer plugin. The following morphological parameters were read out: the number of dendritic branches and the total dendritic branch length (TDBL). Sholl analysis was performed by counting the number of intersections between dendrites and centered on the soma concentric spheres with 10 μm radius increments (78).

#### Analyses of the survival rate

All FOVs including cells and blood vessels were imaged at 14, 25, and 45 dpi under the same settings. The size of each image stack was 635 µm x 635 µm x 200 µm (XYZ). To minimize the effects of cell migration, a safe margin amounting to 100 µm x 100 µm (XY) from the image border was introduced, and cells residing within the margin area were excluded from the analysis. By using blood vessels as a landmark, a cell was considered surviving if its soma was found at the same position in both image stacks (14 and 25 dpi, or 25 and 45 dpi). An offset of ≤ 4 µm was tolerated due to the resolution of the microscope.

#### Analyses of the pCREB expression level and quantification of the Kv1.2 expression

All brain slices were stained and imaged under the same conditions (i.e., antibody concentration, incubation time, laser power, photomultiplier voltage, etc.). Background-subtracted images were generated by estimating background noise from 5 negative control slices, which were stained with secondary antibodies only, and subtracting the median noise value from the original images. Each slice was simultaneously stained with anti-pCREB, anti-GFP (recognizes Twitch-2B), and anti-NeuN antibodies, with secondary antibodies conjugated with Alexa Fluro-488, 594, and 680, respectively. All images were processed using the following protocol in ImageJ: (i) generate 3 substacks for pCREB, GFP, and NeuN staining by splitting the original 3D stack; (ii) draw the ROIs for the adult-born JGCs on the central slice (Z-axis) of the GFP (Twitch-2B) stack; (iii) find the corresponding frame (same depth) in the NeuN stack; (iv) subtract the median background noise value from the NeuN stack; (v) adjust the threshold of the image to highlight all NeuN positive regions and draw ROIs for NeuN-positive cells in the whole field of view; (vi) measure the fluorescence intensity of NeuN-positive mature neurons and GFP-positive adult-born JGCs in the stack of pCREB; (vii) calculate the relative pCREB expression level as the intensity of pCREB fluorescence of each GFP-positive adult-born JGC divided by the median intensity of all NeuN positive mature neurons located in the same field of view.

For quantification of the Kv1.2 expression, the background-subtracted images were generated as described above and a custom-written Matlab code was used to calculate separately the intensity of immunofluorescence in the cell somata and the surrounding neuropil. The relative Kv1.2 expression level was calculated as the ratio of the somatic fluorescence divided by the fluorescence of the surrounding neuropil.

#### Transcriptomic analyses

Read quality of RNA-seq data in fastq files was assessed with ReadQC (ngs-bits version 2019_09). Raw reads were filtered to remove sequencing adapters and for quality trimming using SeqPurge (ngs-bits version 2019_09). Filtered reads were aligned against the reference mouse genome of the Ensembl Mus Musculus GRCm38 using STAR (v2.7.0f), allowing gapped alignments to account for splicing. Low mapping quality reads or those mispairing or multi-mapping were removed using Samtools (v1.9) and visually inspected in the Integrative Genome Viewer (v2.4.19). A matrix of raw counts was built using subread (v1.6.4). Low expressed transcripts were filtered out to minimize the false-positive rate. For each dataset, all transcripts with less than 1 count per million in at least two samples were excluded, leaving 11,611 genes for further differential expression analysis. Differentially expressed genes (DEGs) were identified using edgeR (v.3.24.3) with R (v3.5.2) (https://www.R-project.org/) following the standard workflow. With this method, the size of the library is corrected and differential expression is tested by using negative binomial generalized linear models(79). Genes with an absolute >2-times-fold change between the control and Kv1.2/Kir2.1 groups were considered as DEGs and imported into the online portal Metascape (https://metascape.org/gp/index.html#/main/step1) to run GO biological processes, cell components, molecular functions, and KEGG pathway enrichment analyses (80). In this study, the focus was on signaling pathways downstream of pCREB as well as on the routes leading to CREB phosphorylation.

### Quantification and Statistical Analysis

Statistical analyses were performed using the GraphPad Prism 9 software (GraphPad Software Inc, San Diego, California USA). Shapiro-Wilk test was used to check the normality of data distribution within the data sets. *P* < 0.05 was considered statistically significant. In the case of normally distributed data, the parametric (i.e., Student’s t-test or ANOVA followed by Tukey’s multiple comparison test) otherwise nonparametric (i.e., Mann-Whitney U test or Kruskal-Wallis test followed by Dunn’s multiple comparisons test) tests were used. For Sholl analysis, we used the generalized mixed-effects model with Poisson family (R programming language (81) with lme4 package (82)). P-values were obtained using lmerTest package, which estimates p-values based on Satterthwaite approximation (83). Post-hoc comparisons were done using lsmeans package (84). Box plots were used to present 5 parameters of the dataset: 10th, 25th percentile, median, 75th, and 90th percentile. Unless otherwise indicated, the error bars represent median ± IQR (interquartile range).

